# Predictive Networks Generate Motion-Induced Color Illusions

**DOI:** 10.1101/2025.11.13.688160

**Authors:** Kyohei Ueda, Lana Sinapayen, Eiji Watanabe

## Abstract

Subjective color (SC) refers to chromatic perception induced by achromatic stimuli, as classically demonstrated using Benham’s top. Despite nearly two centuries of investigation, the underlying neural mechanisms remain unclear. We developed artificial neural network (ANN) models trained on natural videos using predictive learning to investigate SC generation. These models successfully reproduced artificial subjective color (ASC) despite the absence of explicit color information in the training stimuli. Critically, ASC characteristics were systematically influenced by the colors of moving objects in the training videos. Using progressively simplified stimuli, ranging from natural scenes to 3D computer graphics to 2D animations, we demonstrated that the colors of moving objects are the primary determinant of ASC properties. We propose that predictive learning establishes color associations with motion patterns, causing the arcs on Benham’s top to be perceived as moving objects with their corresponding colors. These findings suggest that cortical predictive mechanisms may complement retinal processes in generating subjective color phenomena.

## Introduction

Under certain conditions, achromatic visual stimuli can evoke vivid color perception—a phenomenon known as “subjective color“ (SC). This remarkable perceptual experience was first systematically documented by Prevost, who observed the emergence of various colors when white light was presented intermittently (Prevost, 1826). Building upon this foundational work, Fechner demonstrated that colors could also be induced through the rotation of black-and-white patterns, formally coining the term “subjective color” to describe this phenomenon (Fechner, 1838). Despite the various terminologies adopted in subsequent literature, we maintain Fechner’s original nomenclature throughout this paper.

The scientific validity of SC has been established by numerous researchers over nearly two centuries, with Benham’s top (Benham, 1894) emerging as the most widely recognized demonstration of this effect. However, despite extensive investigation (see Cohen & Gordon, 1949; Campenhausen & Schramme, 1995 for comprehensive reviews), the precise neural mechanisms underlying SC remain contentious and incompletely understood.

Figures 1a and 1b illustrate the classic Benham’s top configuration, which consists of a white semicircle and a black semicircle with strategically positioned black arcs inscribed within the white region (black-arc variant). During rotation, observers consistently report chromatic perception along the black arcs. Critically, the phenomenon exhibits strong temporal dependency, with optimal color perception occurring at rotation frequencies of 5–10 Hz. The spatial arrangement of arcs determines the specific colors perceived: arcs positioned at different radial distances from the center evoke distinct colors (position dependency), and these perceived colors systematically reverse when the rotation direction is inverted (direction dependency). In addition to the black-arc variant, a white-arc variant with inverted colors has also been reported (Finnegan and Moore, 1895; Figure 1c).

**Figure 1.**
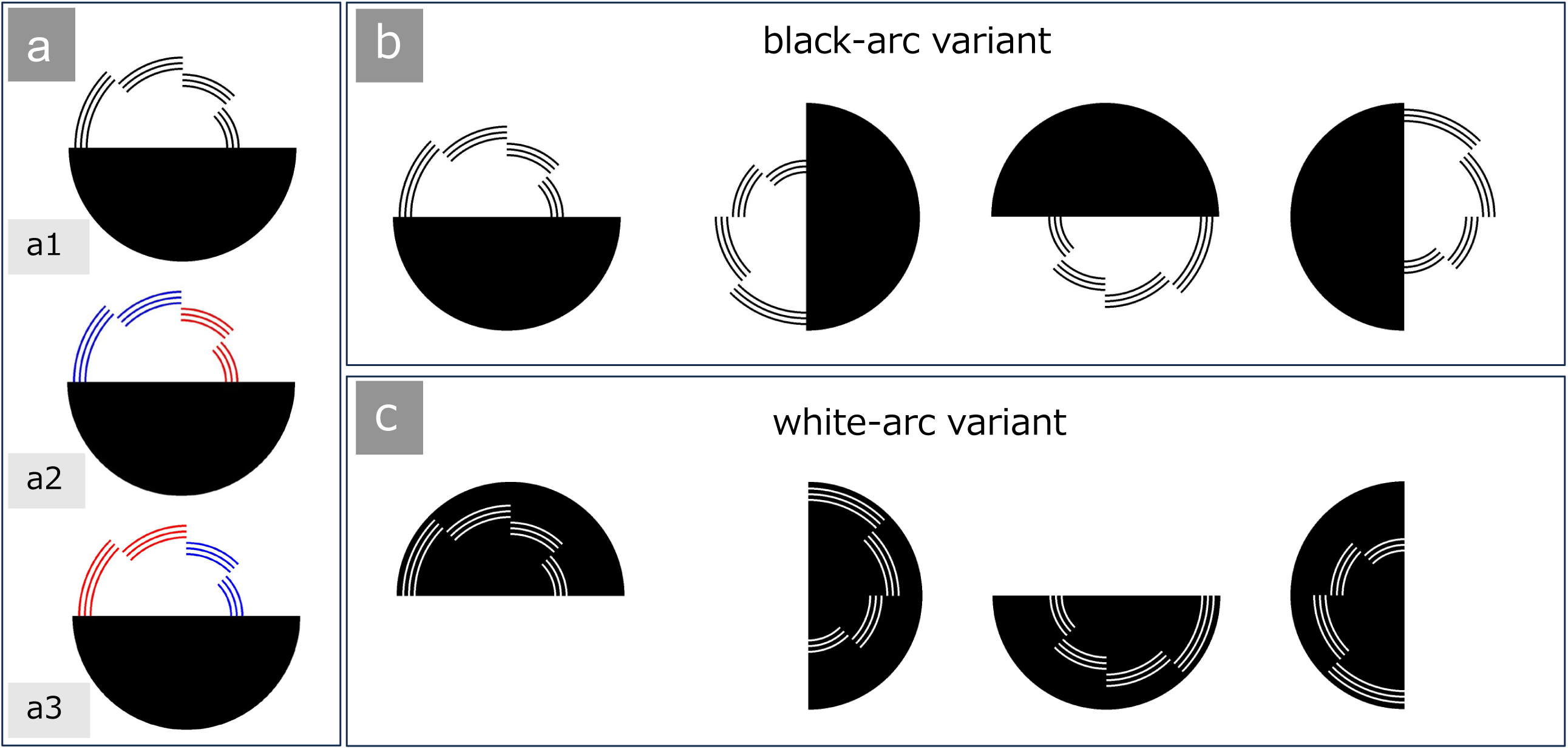
Variations in arc arrangement for Benham’s Top illusion stimuli. (a) Perceived color position depends on rotation direction. (a1) When stationary, the Benham’s top displays uniformly black arcs with no color perception. (a2) Upon rotation, distinct colors are perceived at different radial positions (outer versus inner arcs). (a3) Reversing the rotation direction inverts the order of perceived arc colors and shifts their radial positions. (b) Black arc variant: Stimuli consisting of black arcs on a white background. The stimuli used in this study complete one full rotation across 4 frames. When the four frames are displayed sequentially from left to right, the stimulus rotates counterclockwise; when displayed from right to left, it rotates clockwise. (c) White arc variant: Stimuli consisting of white arcs on a black background. The stimuli used in this study complete one full rotation across 4 frames. When the four frames are displayed sequentially from left to right, the stimulus rotates clockwise; when displayed from right to left, it rotates counterclockwise.

Multiple theoretical frameworks have been proposed to explain SC generation (reviewed in Campenhausen & Schramme, 1995). The first framework emphasizes differential temporal processing of color channels, proposing that flickering black-white transitions induced by rotation generate differential activation patterns across color-processing pathways. The second framework, termed the modulation hypothesis, posits that neural impulse patterns encode specific color information, with rotation-induced flickering modulating these temporal signatures. The third approach emphasizes spatial processing mechanisms analogous to lateral inhibition, focusing on interactions between white background and black arc elements (Campenhausen, 1968; Hasegawa, 1971). Notably, when arc width increases, colors preferentially appear at arc boundaries, suggesting that spatial relationships between pattern elements critically influence SC generation.

While early explanations focused primarily on retinal processing, accumulating evidence points toward cortical involvement. Individual differences in perceived color categories cannot be adequately explained by cone cell characteristics alone (Knehr, 1953). Furthermore, asymmetric outer arcs influence inner arc color perception even when inner arcs remain symmetric across rotation directions (Le Rohellec et al., 1992)—an effect that exceeds the spatial scale of retinal circuits and implicates higher-level cortical processing. Functional magnetic resonance imaging studies have provided direct evidence for cortical feedback mechanisms in SC perception (Tanabe et al., 2011). Collectively, these findings suggest that SC emerges from hierarchical processing involving both initial retinal computations and subsequent cortical integration.

To elucidate the neural origins of SC perception, we developed computational models using artificial neural networks (ANNs). Since their inception as neuron-inspired computational frameworks (McCulloch and Pitts, 1943), ANNs have proven invaluable for modeling diverse brain functions, particularly visual processing. The success of ANN modeling of the ventral visual stream has been remarkable, with architectures inspired by Hubel and Wiesel’s cascade hypothesis (Hubel and Wiesel, 1962)—from the Neocognitron (Fukushima, 1980) to convolutional neural networks such as LeNet-5 (LeCun et al., 1998) and AlexNet (Krizhevsky et al., 2012)—consistently reproducing neurophysiological findings (Richards et al., 2019). Research on dorsal stream models, which process spatiotemporal information such as object motion, is also advancing rapidly (Mineault et al., 2021; Sarch et al., 2023; Choi et al., 2023).

Beyond unidirectional processing models, research on bidirectional information flow in the cerebral cortex (Gilbert and Li, 2013) is also progressing. A prominent example is predictive coding (Rao and Ballard, 1999), which posits that higher-order cortex contains an internal model of the world in which predictions are matched against inputs at lower cortical levels, and the resulting prediction error is utilized for learning (see Figure 2a). This concept evolved from early work by Kawato and colleagues (Kawato et al., 1993) into frameworks such as predictive coding (Rao and Ballard, 1999) and the free-energy principle (Friston, 2010). Within this context, we previously proposed that visual illusions arise through top-down predictive signals (Watanabe et al., 2010).

**Figure 2.**
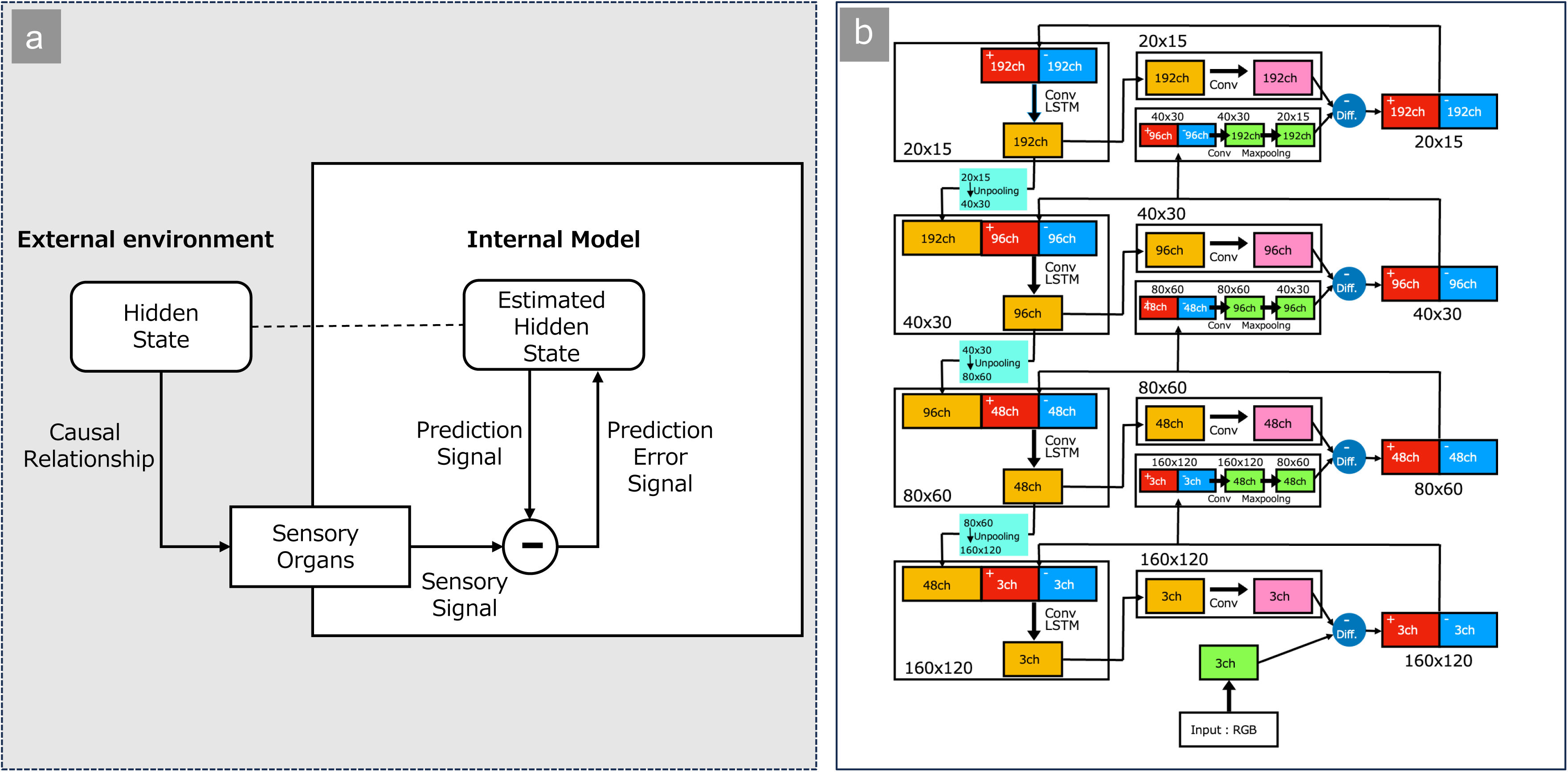
Predictive coding framework and its implementation in PredNet. Left: Schematic illustration of the predictive coding framework. The external environment generates hidden states that causally produce sensory input through sensory organs. An internal model estimates these hidden states and generates prediction signals. These predictions are compared with incoming sensory signals, and the resulting prediction errors are fed back to update the internal model. Right: Implementation of the predictive coding principle in the PredNet deep learning architecture. The network is organized hierarchically across multiple spatial scales (ranging from 160×120 to 20×15 pixels). Each layer contains convolutional long short-term memory (LSTM) units that generate predictions, which are then compared with incoming sensory features. Prediction errors (denoted as “Diff”) are propagated forward through the hierarchy, while prediction signals are fed back to refine internal representations. This hierarchical, error-driven architecture mirrors the predictive coding framework, enabling both temporal prediction and representation learning from visual input.

Predictive coding has been implemented as an ANN through PredNet (Lotter et al., 2017; Figure 2b), which has demonstrated particular success in reproducing motion illusions (Watanabe et al., 2018; Kobayashi et al., 2022; Kobayashi and Watanabe 2023) and the flash-lag effect (Lotter et al., 2020), and can even generate novel optical illusions when combined with genetic algorithms (Sinapayen and Watanabe, 2021). Furthermore, correlations with cortical neurons responding to subjective contour stimuli have been identified (Lotter et al., 2020). While PredNet does not necessarily exhibit one-to-one correspondence with physiological structures in the manner of CNNs modeling the ventral stream, it represents a promising framework for validating cortical information processing principles through modeling of subjective visual phenomena.

In this study, we developed predictive network models capable of generating artificial subjective color (ASC) and leveraged these computational frameworks to investigate the underlying mechanisms of SC perception.

## Results

### Human Psychological Experiments

To validate our experimental paradigm, we first conducted online psychological experiments to characterize human perceptual responses to the Benham’s top stimulus used in our ANN studies. We employed a Benham’s top that completed one full rotation in four frames (Figure 1b and 1c, supplementary videos). This discrete rotation pattern elicits only two distinct subjective colors, facilitating quantitative analysis.

Given the limited literature on white-arc variants of Benham’s top, we deemed this validation essential. While Finnegan and Moore first reported color perception with white-arc variants in 1895, subsequent studies have yielded conflicting results—some denying color perception entirely (Piéron, 1922) while others supporting it (Mitsuhiro and Kitaoka, 2024). Moreover, phenomena such as hue pattern reversal and rotation direction dependence have not been systematically documented. Therefore, we conducted an online survey in which 98 participants indicated perceived colors on a standardized color map.

The results are shown in Figure 3 (rose plot) and Table 1 (HSV data). For the black-arc variant with counterclockwise (ccw) rotation, warm colors (hue angle 53.3° ± 73.4°, mean ± SD) were perceived on inner arcs and cool colors (260.3° ± 63.0°) on outer arcs (included angle between the two colors: 153.0°), whereas clockwise (cw) rotation reversed the hue positions (63.2° ± 68.7°; 268.5° ± 57.5°; included angle: 154.7°). The white-arc variant showed opposite hue positioning relative to the black-arc variant (see Table 1 for details, “arc-color dependency”). Illusory colors of white-arc variants were more difficult to perceive—12.2% of participants reported no color perception (HSV = 0,0,0) for inner arcs and 14.3% for outer arcs, compared to 0% for both positions in black-arc variants. Under all conditions, saturation exceeded 20%, the threshold at which colors are considered clearly visible (Sharma and Dubey, 2012).

**Figure 3.**
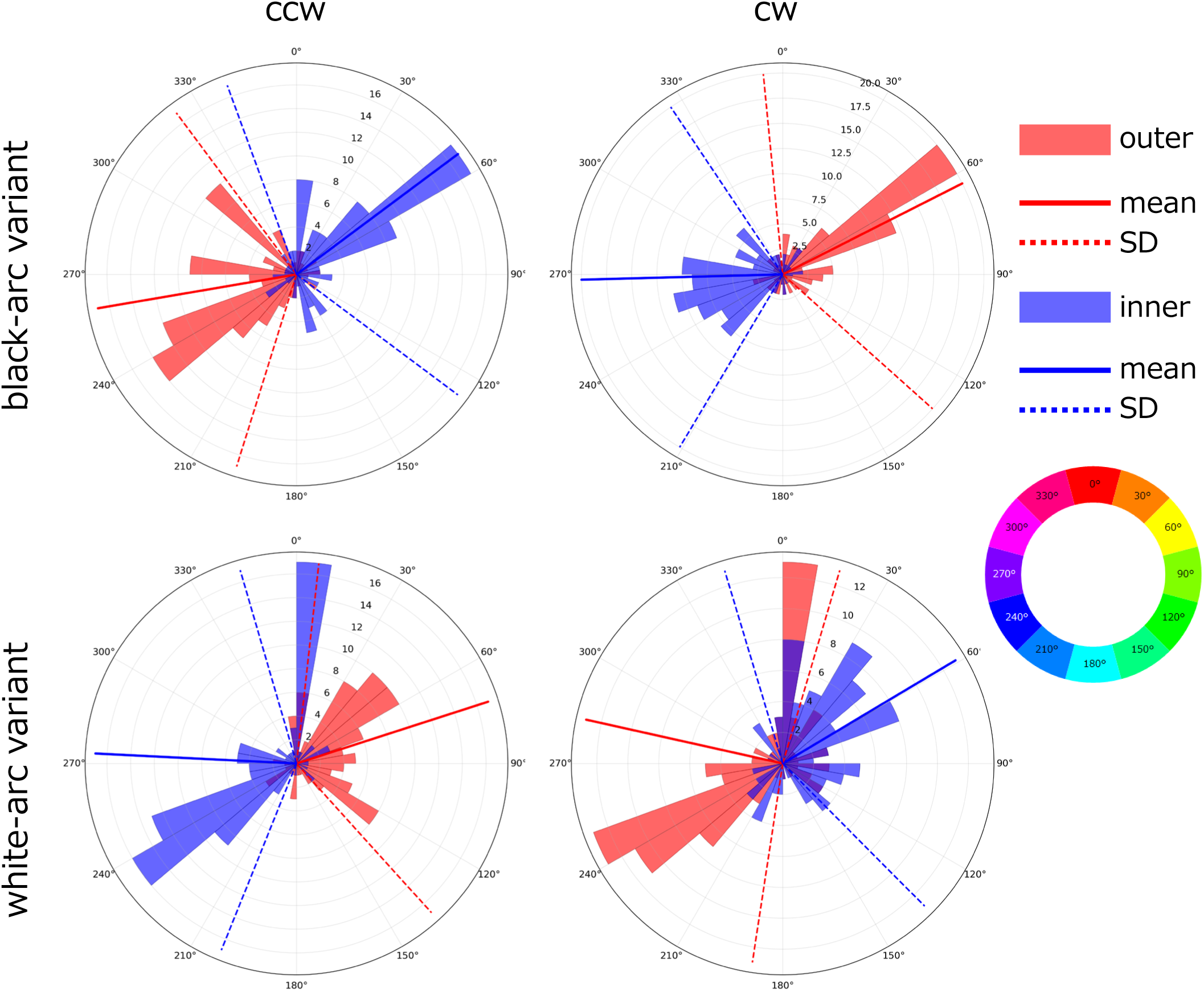
Rose plots of perceived hues for Benham’s top stimuli. Polar histograms show the distribution of reported hues perceived by participants (n = 98) as a function of angular position (hue angle in degrees). The radial extent of each bin represents the number of observers reporting each hue category. Results are presented separately for the black-arc variant (top row) and white-arc variant (bottom row) under counterclockwise (ccw, left column) and clockwise (cw, right column) rotation conditions. Red histograms represent hues perceived at the outer arc; blue histograms represent hues perceived at the inner arc. Solid lines indicate the mean hue angle, and dotted lines indicate ±1 standard deviation (SD) from the mean. The color wheel (right panel) provides a reference for the correspondence between angular position and hue value.

In the psychological experiment, we confirmed “position dependency” and “direction dependency,” and additionally discovered “arc-color dependency.” In subsequent computational modeling experiments, we evaluated the model’s validity primarily based on these three dependencies.

### ANN Models Trained on First-Person View Natural Scene Videos

We subsequently employed artificial neural networks (ANNs) trained on first-person view natural scene videos to implement predictive learning and analyzed their responses to Benham’s top stimuli. Critically, during training, the ANNs were never exposed to Benham’s top stimuli nor provided with any indication that such stimuli should evoke colors—we treated the networks as naive observers mimicking human visual experience.

We trained an ANN using 17 randomly selected first-person view natural scene videos, and a representative example of the results is shown in Figure 4 (rose plot) and Table 2 (HSV data). When a rotating Benham’s top was input to the trained ANN, pale colors appeared along the arcs of the predicted image. The colors exhibited the three dependencies observed in human subjects: position dependency, direction dependency, and arc-color dependency. When a stationary Benham’s top was input, almost no color was observed along either the black arcs (saturation: 0.7 ± 0.6 for outer arc, 0.6 ± 0.6 for inner arc) or white arcs (2.7 ± 0.8, 1.7 ± 0.6). Because the ANN model reproduced at least these three dependencies observed in human subjective color perception, we designated this phenomenon “Artificial Subjective Color (ASC).”

**Figure 4.**
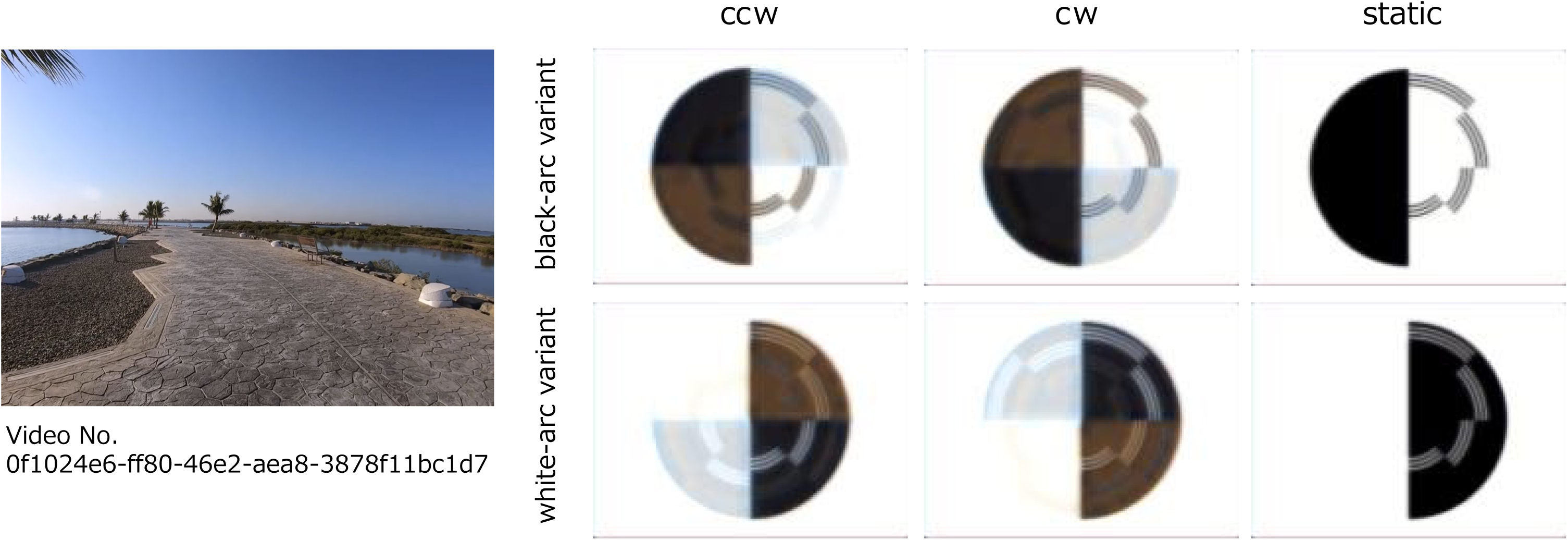
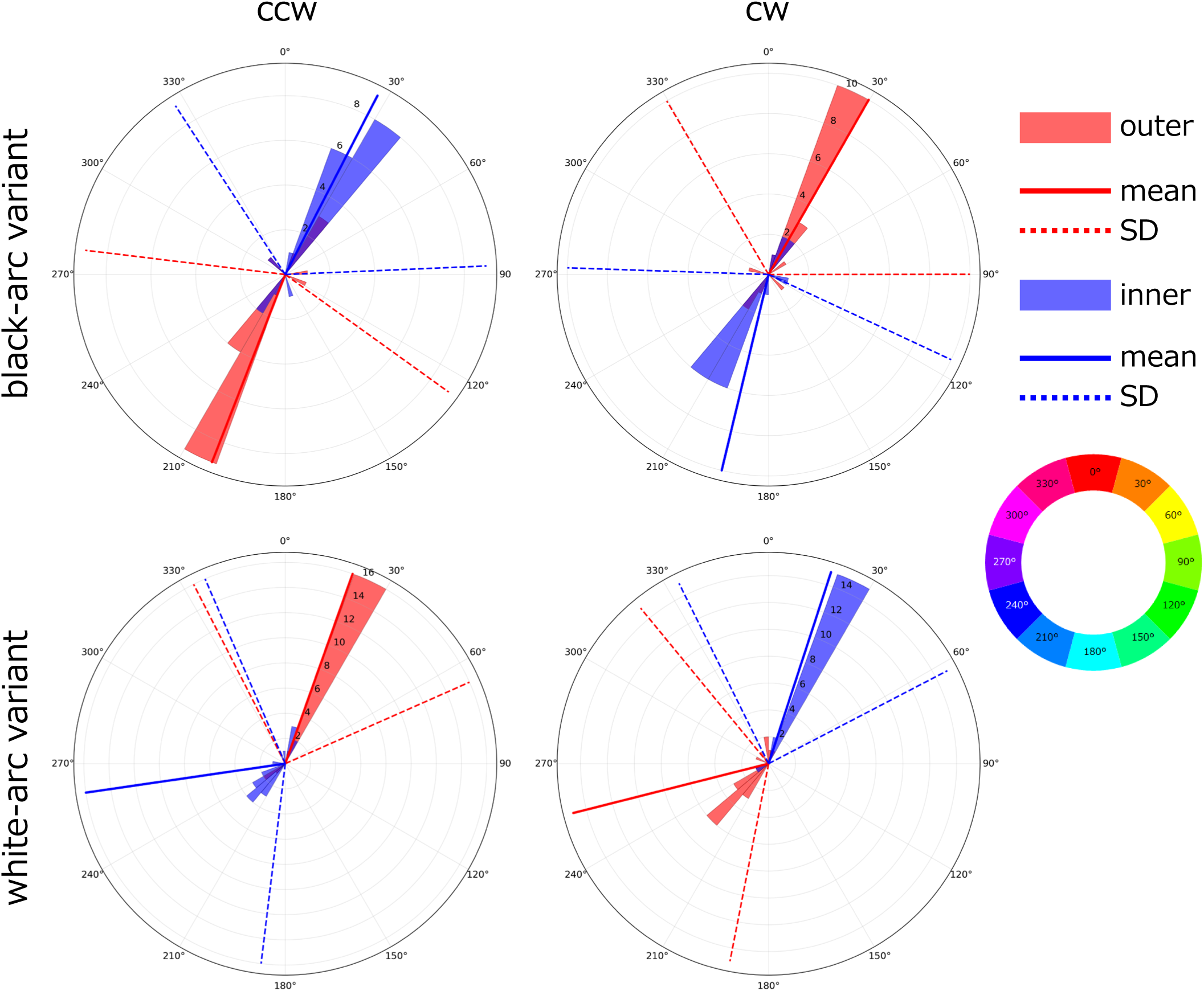
PredNet predictions for Benham’s top variants after training on first-person natural scenes (Ego4D dataset). (4a) *Left:* An example frame from the Ego4D training video (video ID: 0f1024e6-ff80-46e2-aea8-3878f11bc1d7) depicting a first-person view of two individuals cycling through an urban environment. Although most of the actual video footage captures people riding bicycles together, an image that does not show individuals was used in order to protect personal information. The first 40,000 frames of each video were used for training. *Right:* Representative predicted images generated during the inference phase when Benham’s top stimuli were presented as novel inputs. Two stimulus variants were tested, featuring either black (top row) or white (bottom row) arc segments. Under rotation (counterclockwise, ccw; clockwise, cw), the model’s predictions exhibited distinct color patterns at the inner and outer arc positions, with their relative spatial arrangement depending on the direction of rotation. By contrast, no color percepts were observed under static conditions. (4b) Rose plots depicting the distribution of predicted hues from the neural network model (n = 20 trials). Results are shown for the black-arc variant (top row) and white-arc variant (bottom row) under counterclockwise (ccw, left) and clockwise (cw, right) rotation conditions. Color coding follows the convention established in Figure 2: red histograms represent hues predicted at the outer arc position, and blue histograms represent hues at the inner arc position. Solid lines indicate the mean direction, and dotted lines indicate ±1 standard deviation.

However, several differences from the human data also became apparent. First, saturation was lower (minimum 7.5, maximum 25.4) compared to human data, with only pale colors observed (Figure 4a). Furthermore, while this model expressed hues (27.3° ± 60.3° for inner black-arc ccw, 201.3° ± 75.6° for outer black-arc CCW, 193.5° ± 78.5° for inner black-arc cw, 29.8° ± 60.2° for outer black-arc cw; included angles: 174.0° and 163.7°) similar to those perceived by human subjects, numerous models obtained from other training videos expressed hues that deviated from the perceived hues (see Figure S1). In two training videos, no clear ASC occurred (Figure S1i and S1m). These results suggest that while ASC arises from learning on first-person view natural scenes, its specific characteristics depend on the training content.

### ANN Trained on 3D Computer Graphics Videos

To systematically investigate which video features influence ASC generation, we created controlled 3D computer graphics (3DCG) training videos depicting “a walking person in a town” (Figure 5a, supplementary video). These videos reproduced key elements of natural footage—colors, shapes, backgrounds, objects, shadows, and motion dynamics—while excluding potentially confounding factors such as camera shake and blur effects (see Methods for details).

**Figure 5.**
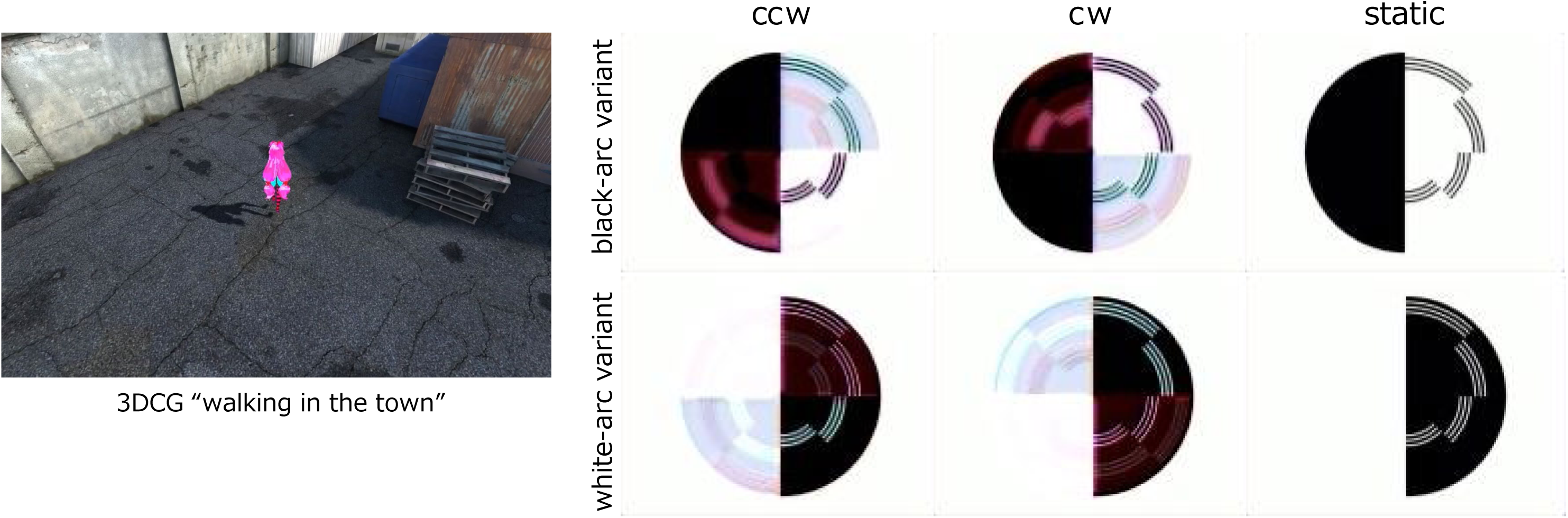
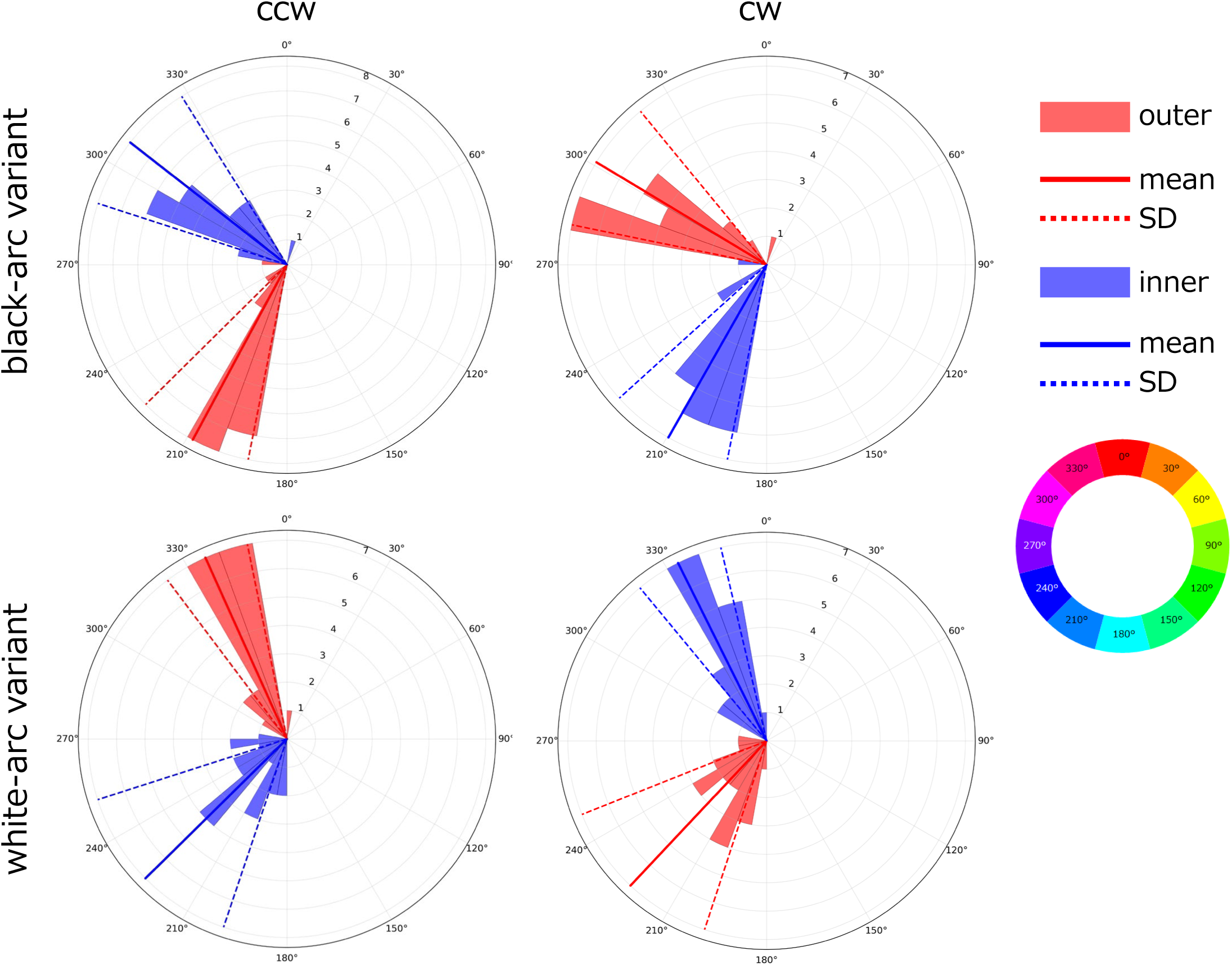
PredNet predictions for Benham’s top variants after training on artificial 3D scenes. (5a) *Left:* Example frame from a computer-generated 3D video showing a human character walking through a town environment, used during PredNet training. *Right:* Representative predicted images obtained during the inference phase when Benham’s tops were presented as novel inputs. As in Figure 4a, two stimulus variants were tested (black-arc, top row; white-arc, bottom row) under counterclockwise (ccw), clockwise (cw), or static conditions. Compared with the model trained on natural scenes (Figure 4a), the illusory colors predicted for the rotating conditions appear more vivid and distinct, whereas static inputs again elicited no color. These results highlight the influence of training data statistics—in this case, structured artificial 3D motion—on the emergence and clarity of motion-contingent color illusions in the model’s predictions. (5b) Rose plots of predicted hues generated by the neural network model (*n* = 20). Results are shown for the black-arc variant (top row) and the white-arc variant (bottom row) under counterclockwise (ccw, left) and clockwise (cw, right) rotation conditions. Blue and red color coding follows the same convention as in Figure 4b.

Training on 3DCG videos again produced clear ASC responses to rotating Benham’s top (Figure 5, Table 3), indicating that the rich complexity of natural videos is not required for ASC generation. For the black-arc variant with counterclockwise (ccw) rotation, warm colors (hue angle 308.0° ± 20.0°, mean ± SD) were perceived on inner arcs and cool colors (208.3° ± 17.0°) on outer arcs, whereas clockwise (cw) rotation reversed the hue positions (209.6° ± 18.2°; 301.1° ± 19.5°). The white-arc variant showed opposite hue positioning relative to the black-arc variant (see Table 3 for details). This approach produced more vivid colors than the ANN trained on first-person view natural scenes (mean saturation minimum 11.3, maximum 34.1). The resulting ASC exhibited approximately 100-degree hue differences between the two colors. When a stationary Benham’s top was input, almost no color was observed, consistent with natural scene video results (Figure 5a; see saturation values for Ba st and Wa st in Table 3).

### ANN Trained on Simple 2D Computer Graphics Videos

To enable more precise quantitative analysis, we further simplified the training stimulus to mathematically defined 2D graphics. Focusing on two elements—the background and moving objects—we replaced walking figures by colored squares (red, blue, or green), constrained movement to 2D uniform linear motion, created backgrounds from randomly colored dots, and eliminated camera movement entirely (Figure 6a, supplementary videos).

**Figure 6.**
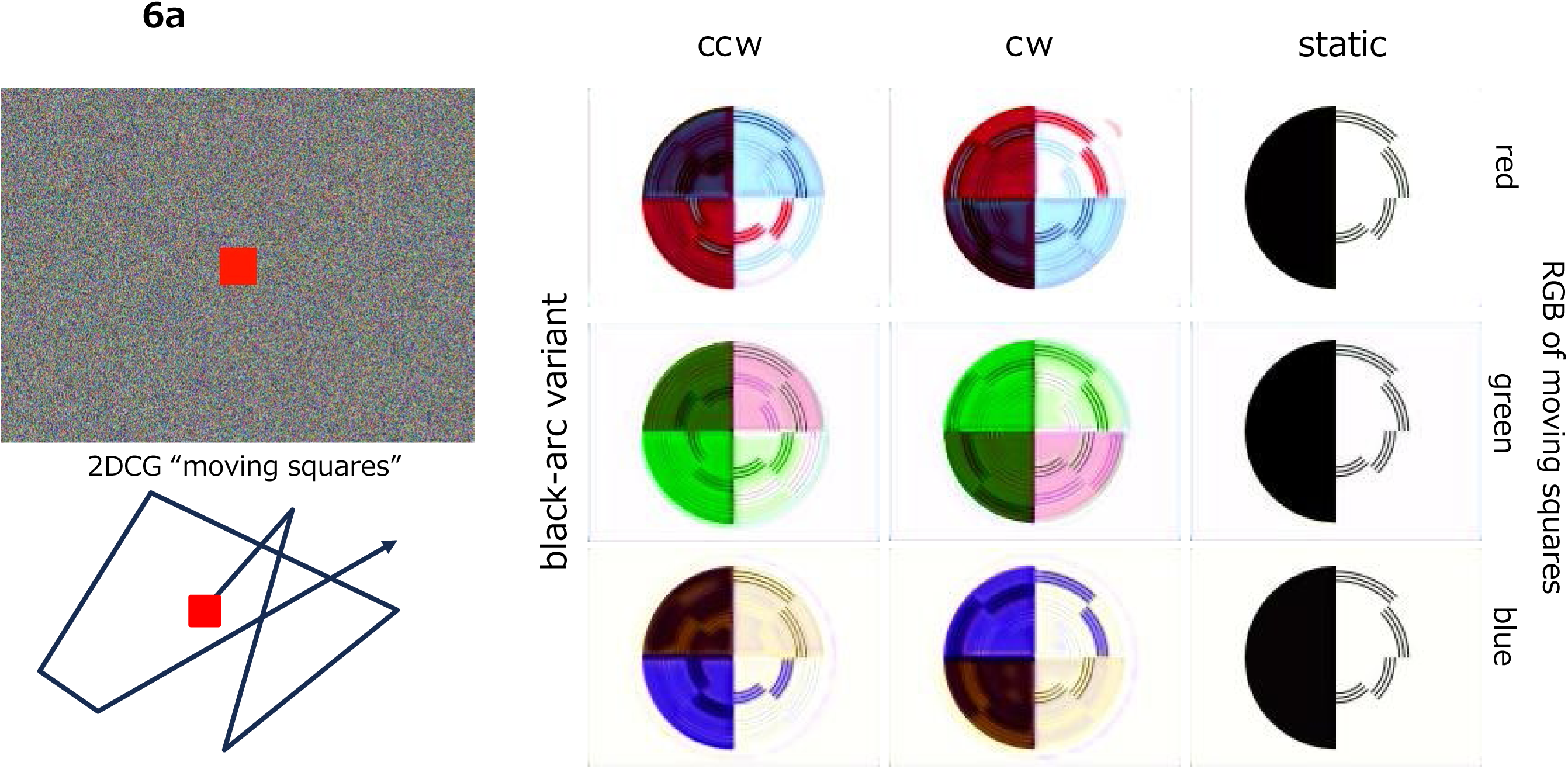
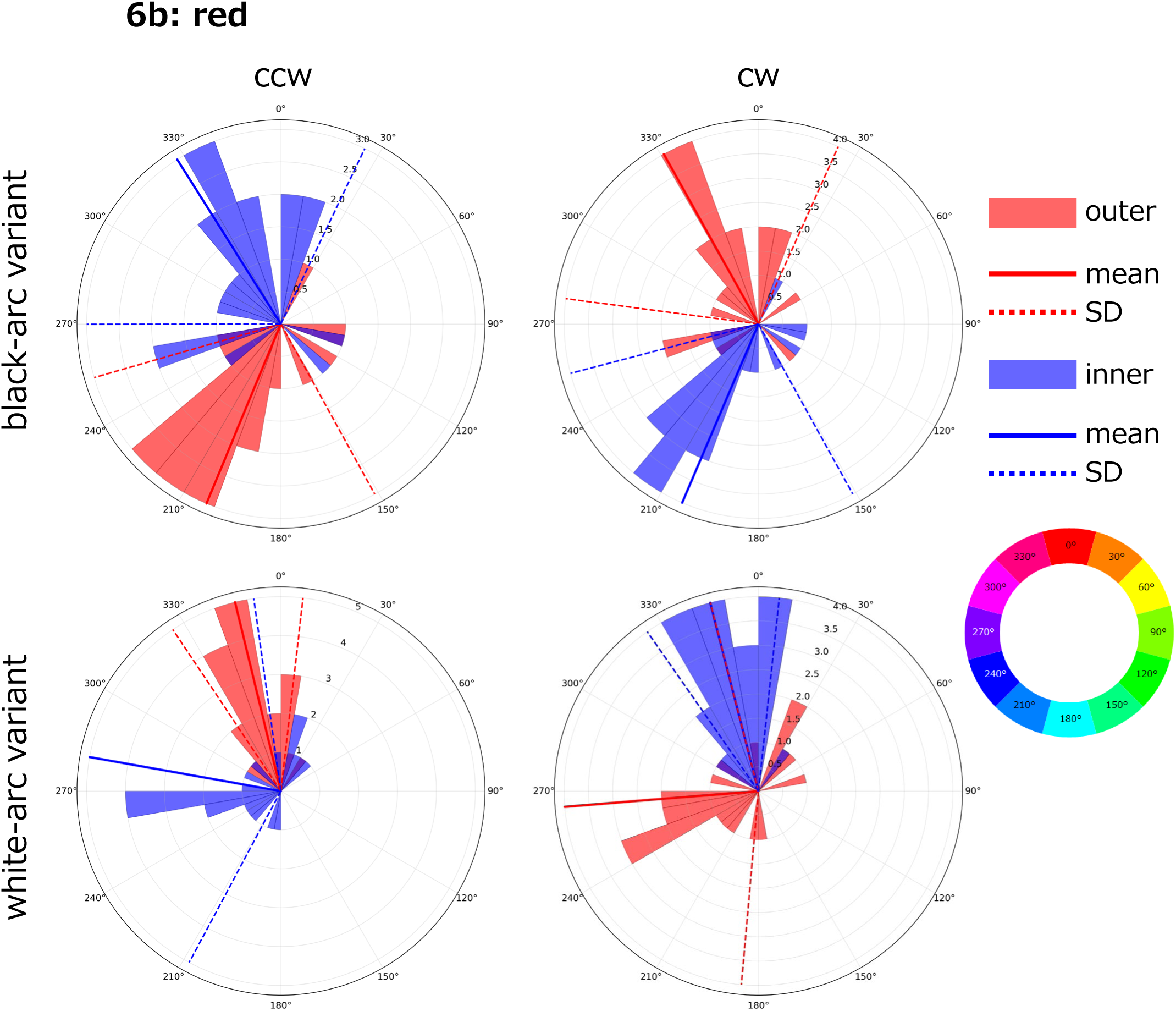
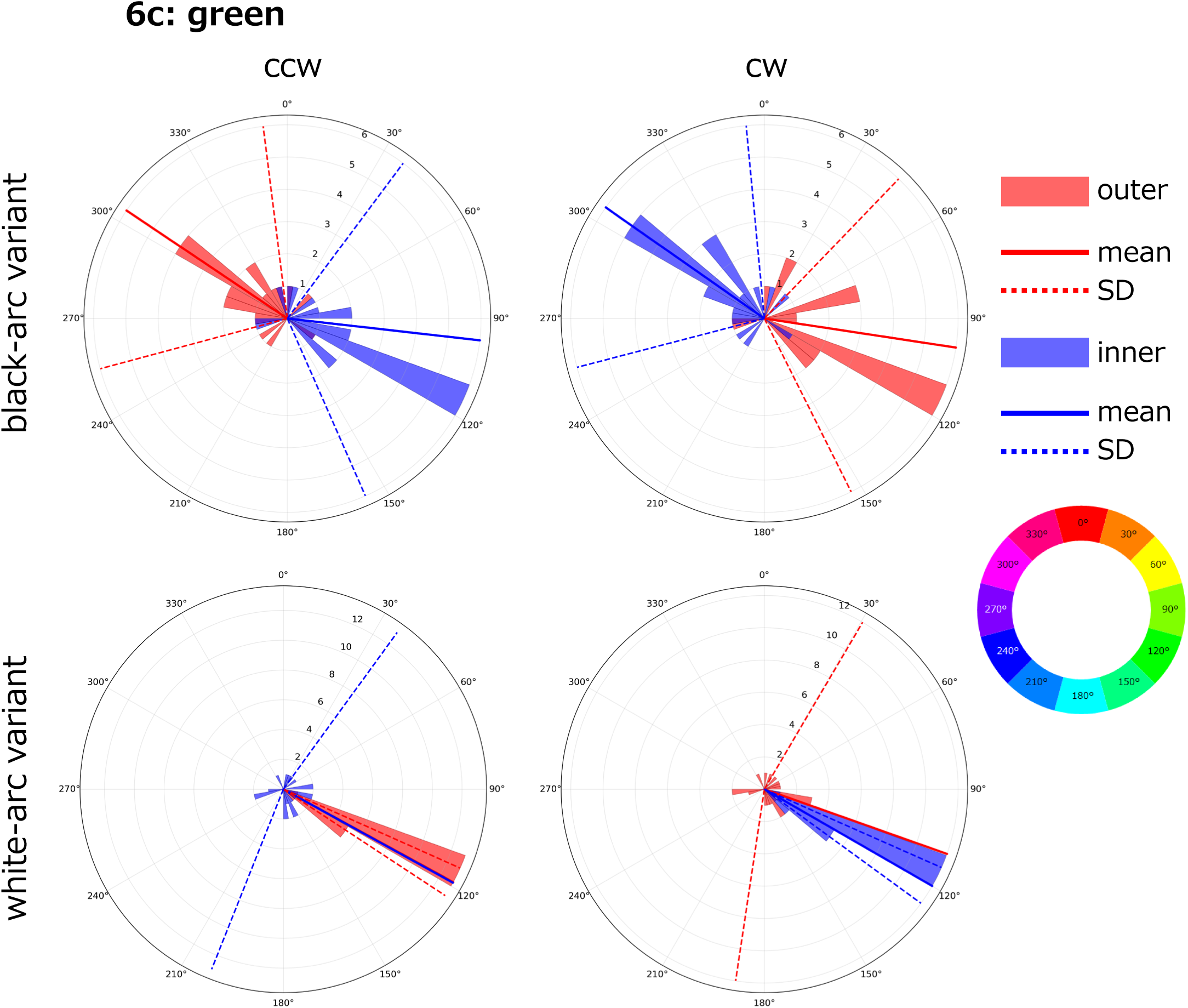
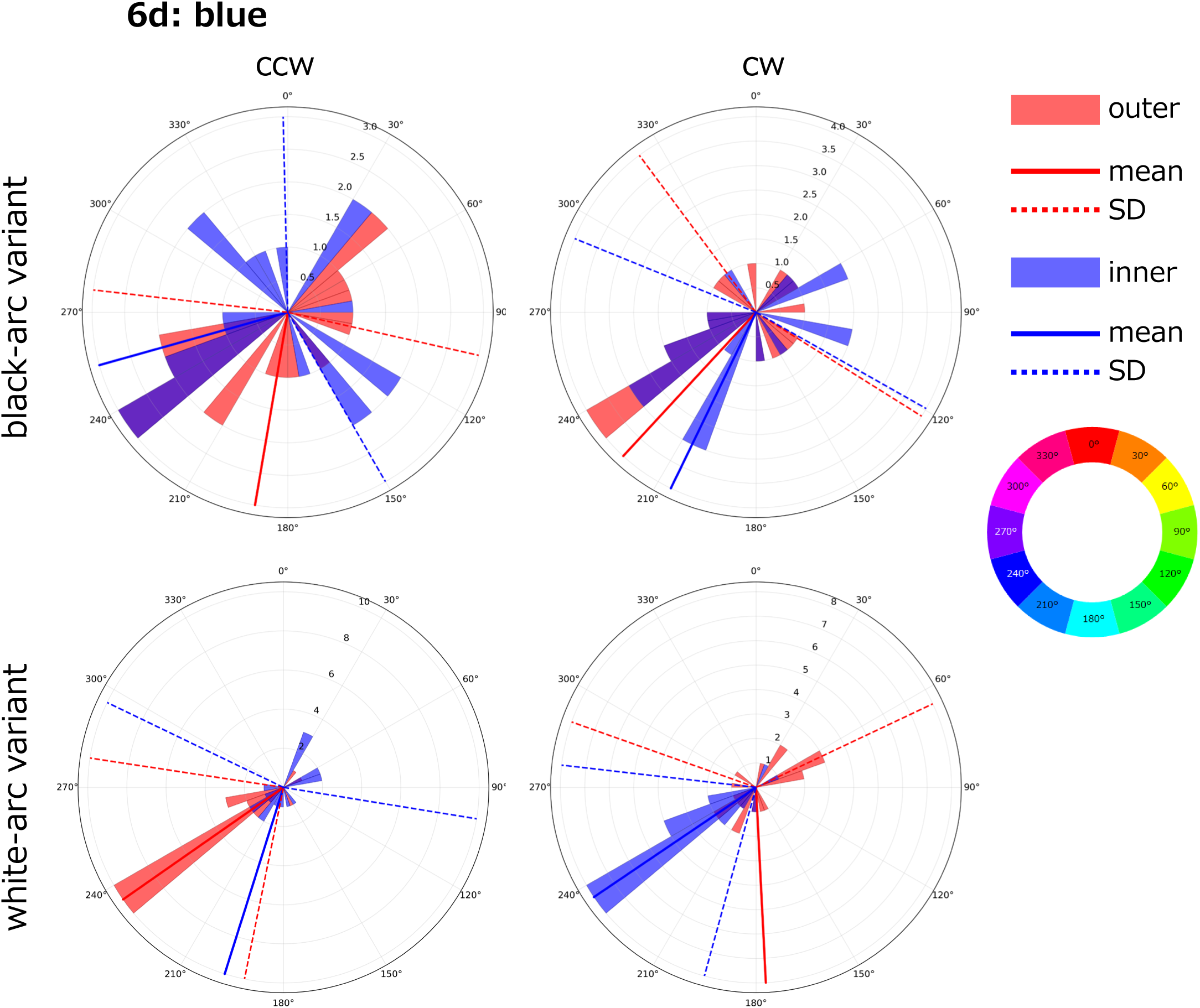
PredNet predictions following training on simplified 2D motion stimuli. (6a) *Left:* Example frame from training videos showing a colored square (red, green, or blue) moving along a random trajectory against a background of dynamic colored noise (2D computer graphics). *Right:* Representative predicted images generated during inference when Benham’s top was presented as novel inputs (black-arc variant). Compared with models trained on natural scenes (Figure 4a) or 3D artificial scenes (Figure 5a), the predicted illusory colors are notably more vivid. Furthermore, the colors appearing in the arcs tend to match those of the moving square used during training, indicating that the model’s predictions reflect the dominant motion–color contingencies experienced during training. Consistent with previous experiments, no illusory colors emerge under static conditions. (6b, 6c, 6d) Rose plots of predicted hues generated by the neural network model (n = 20). In 6b, the network was trained using moving red objects; in 6c, using moving green objects; and in 6d, using moving blue objects. Results are shown for the black-arc variant (top row) and white-arc variant (bottom row) under counterclockwise (ccw, left) and clockwise (cw, right) rotation conditions. Color coding follows the convention established in Figure 4a.

Using the red-square video, ASC was successfully reproduced even with these highly simplified stimuli (Figure 6a, 6b, and Table 4a). For the black-arc variant with counterclockwise (ccw) rotation, warm colors (hue angle 327.7° ± 57.8°, mean ± SD) were perceived on inner arcs and cool colors (202.6° ± 51.4°) on outer arcs, whereas clockwise (cw) rotation reversed the hue positions (203.1° ± 52.1°; 330.9° ± 53.4°). The white-arc variant showed opposite hue positioning relative to the black-arc variant (see Table 4a for details). These results suggest that one arc color of the Benham’s top is influenced by the color of the moving object. From this perspective, the same principle is likely applicable to 3DCG experiments. Coincidentally, the hue of the walking figure resembles the hue appearing on one of the arcs (inner or outer) predicted by ANN models.

Similar results were observed in experiments using blue and green videos (Figure 6a, 6c, 6d and Tables 4b, 4c). However, only one color was observed in the white-arc Benham’s top experiments, not two. Furthermore, in experiments using blue videos, the learning outcomes for the black-arc Benham’s top showed significant variation.

One arc color was strongly influenced by the colors of moving squares in the training videos, indicating that moving object colors were the primary determinant of ASC characteristics. These results demonstrate that ASC generation depends fundamentally on color biases acquired during learning, particularly those associated with moving objects. However, the high variance in learning outcomes (especially for blue stimuli) suggests that 2D stimuli may provide insufficient information to fully capture the complexity of Benham’s top color perception.

## Discussion

We successfully developed artificial neural network (ANN) models that reproduce subjective color (SC) phenomena. The artificially generated subjective colors (ASC) depend on the characteristics of training videos, with the colors of moving objects exerting a particularly strong influence on the resulting color perception. Our findings establish that ASC emerges from predictive learning on visual sequences and is systematically influenced by the color statistics of moving objects in training data, providing a computational framework for understanding subjective color phenomena.

### Assumed Mechanism of ASC Generation

We propose the following mechanism for ASC generation based on predictive learning principles. Objects with rapidly changing positions represent particularly challenging prediction targets. Consequently, ANNs trained for predictive tasks are disproportionately influenced by moving objects during the learning process. When training videos contain systematic color biases in moving objects, ANN models develop corresponding color biases in their predictions of motion-related features. This framework provides a clear explanation for why colors appear on Benham’s top arcs when they are conceptualized as moving objects.

Analysis of internal model activity during training with computer graphics reveals that the black semicircle of the top appears to move rotationally together with the arcs (Figure S2). The relative positioning of arcs—whether inner or outer—alternates depending on rotation direction. This pattern becomes interpretable when we consider the black semicircle and its adjacent arcs as constituting “one coherent moving object.” Arcs moving in the same direction as the rotation become integrated into this moving object representation, while counter-rotating arcs are processed as background elements. This conceptual framework naturally accounts for the direction-dependent differences in color appearance observed in SC phenomena. Regarding positional dependency, the internal activation patterns also provide insights. Activity in blocks associated with the outer arc (orange boxes) and inner arc (blue boxes) is processed separately, and their activation phases are inverted with respect to excitation and inhibition. This indicates that positional dependency is expressed through internal processing.

Furthermore, these blocks appear to be related not only to the arcs but also to various static elements such as the disk shape and disk center. Indeed, complementary colors in ASC appear not only on the arcs but also in the background area, suggesting a complex interaction between the moving object and its surrounding context. This interaction is considered important for understanding the positional dependence of black and white contrast observed in SC phenomena. When dynamic and static elements function as complementary components within the visual system, their interaction becomes a fundamental determinant of color perception. In this respect, our ANN model can reproduce a mechanism similar to the spatial interaction hypothesis discussed in the introduction.

### Implications for SC Generation Mechanisms

Our findings also invite reconsideration of the role of central information processing in subjective color generation. Sadato’s research group demonstrated through fMRI analysis that feedback signals from the cerebral cortex contribute to Fechner color perception (Tanabe et al., 2011). The cerebral cortex not only extracts features from peripheral sensory signals but also engages in extensive feedback processing. While the precise role of this feedback varies across contexts, it is thought to convey information related to attention, expectation, perceptual tasks, working memory, and motor commands (Gilbert and Li, 2013). Within predictive coding frameworks, such feedback signals are theorized to represent predictive signals themselves (Rao and Ballard, 1999). The finding that ASC emerges from ANN prediction signals in our study provides compelling support for this theoretical perspective.

Although ANN experiments permit controlled long-term learning paradigms, conducting equivalent experiments with human participants remains impractical. Therefore, direct verification of the relationship between moving object processing and color perception in humans presents significant methodological challenges. However, indirect evidence might be obtained by combining large-scale surveys of subjective color perception across populations living in diverse color environments with quantitative measurements of those environmental color statistics. Recent research demonstrates that perception of geometric illusions differs markedly between individuals from urban versus rural environments (Kroupin et al., 2025), suggesting that environmental experience shapes fundamental perceptual processes.

While the hypothesis that Benham’s top colors result from experiential learning processes may initially appear speculative, approximately 200 years have elapsed since the initial discovery of subjective color phenomena (Prevost, 1826). This accumulated body of evidence now warrants exploration of novel theoretical approaches to understanding these enduring perceptual mysteries.

## Methods

### Psychological Experiments

#### Participants

Participants were recruited through Prolific Academic (https://www.prolific.co/). The total sample comprised 98 individuals (50 males, 47 females, 1 unspecified; mean age = 34.64 years, SD = 10.41). Participants represented 24 countries, with no geographical restrictions imposed. Compensation was provided at an hourly rate of £12.00. Eligibility criteria included normal visual acuity and normal color vision, and participants completed the experiment on personal computers. Screen size, resolution, and calibration were not standardized. Participants used Chrome, Firefox, or Safari browsers. The experimental procedure received approval from the Life Science Research Ethics Committee of the National Institutes of Natural Sciences (Project ID: EC01-078), and informed consent was obtained from all participants.

#### Procedure and Stimuli

The experiment was developed and hosted online using Gorilla Experiment Builder (www.gorilla.sc) (Anwyl-Irvine et al., 2020). The procedure consisted of six sequential stages: (1) informed consent, (2) experimental overview, (3) participant information collection, (4) Benham’s top explanation, (5) practice session, and (6) main experiment.

After providing informed consent, participants completed a questionnaire assessing eye color, living environment, visual acuity, and vision characteristics. We then presented illustrations of the Benham’s top stimuli, explaining the distinction between inner and outer lines (Figure 1a). Subjects were asked to describe the colors appearing in the two outer sets of arcs and the two inner sets of arcs. Participants were instructed to use full-screen mode throughout the experiment.

The practice session included three tasks: (1) stationary Benham’s top, (2) stationary colored Benham’s top, and (3) rotating Benham’s top with colored inner and outer lines. This session confirmed proper video functionality and familiarized participants with the color-matching procedure using mouse-controlled adjustments of saturation, hue, and brightness.

The main experiment comprised 16 trials. Participants reported perceived colors for inner and outer lines across eight stimulus conditions: two Benham’s top types (standard and black-white inverted) × two rotation speeds (20 fps and 30 fps) × two rotation directions (clockwise and counterclockwise) (https://doi.org/10.6084/m9.figshare.30215359). Benham’s top animations consisted of four images representing 90° rotational increments, with quarter-arc sections designed to elicit perception of distinct colors in inner and outer line regions.

Stimulus presentation order was randomized across participants. Response time ranged from 15 to 30 seconds per stimulus, with a total experiment duration of 10 to 15 minutes. Task materials are available on Gorilla Experiment Builder (https://app.gorilla.sc/openmaterials/1002694). Stimulus images were generated using custom Python scripts. Responses were recorded as RGB values (0–255) and subsequently converted to HSV color space for consistency with neural network model analyses.

## Model Experiments

### Deep Neural Networks

We implemented PredNet (Lotter et al., 2016) in PyTorch, adapting the original Keras implementation (https://doi.org/10.6084/m9.figshare.30215359). This represents a new PyTorch implementation, distinct from the previously used Chainer version (Watanabe et al., 2018) due to that library’s discontinuation. Following the hyperparameter configurations from prior natural image sequence learning studies (Lotter et al., 2016), we employed a 4-layer architecture with 3×3 convolutional filters and channel dimensions of 3, 48, 96, and 192 for successive layers. Model optimization utilized the Adam algorithm with default parameters. Input images were standardized to 160×120 pixels in RGB format.

#### Training

Training utilized the video datasets described below. Videos were decomposed into sequential frames and input chronologically to the network. PredNet generates predicted images based on previous video frames, with current frames serving as ground truth for learning. This creates an autoencoder-like architecture for future frame prediction, with Long Short-Term Memory (LSTM) units integrated into convolutional processing to maintain temporal information.

The loss function computed the mean squared error between ground truth and predicted images at the first layer. Batch size was set to 20 images, with backpropagation-based parameter updates after each batch. Models were sampled at fixed intervals during training—not to minimize loss but to identify models producing illusory color phenomena. To ensure training stability, we used predetermined initial values confirmed to reduce loss in preliminary experiments, as training success was highly sensitive to initialization. Training and inference were performed using NVIDIA RTX A6000 GPUs.

### Training Datasets

ANNs were trained on three video dataset types:

1. Ego4D (https://ego4d-data.org/): To approximate human visual experience, previous studies used the First-Person Social Interactions Dataset (Watanabe et al., 2018). However, as these video data are no longer publicly available, we utilized the newly developed Ego4D dataset. Ego4D is a large-scale, first-person view dataset and benchmark suite collected across 74 worldwide locations in 9 countries, containing over 3,670 hours of daily-life activity video. Twenty videos featuring camera and subject movement were randomly selected, and three short videos were excluded. The identification numbers of the selected videos are indicated in Figure 4a and Figure S1.
2. 3D Computer Graphics (https://doi.org/10.6084/m9.figshare.30215359): Artificial animations created using Unity software, featuring simple townscape environments with pedestrians. Camera movement tracked subjects while maintaining varied positioning rather than fixed relative positions.
3. 2D Computer Graphics (https://doi.org/10.6084/m9.figshare.30215359): Animations featuring three colored squares moving in 2D space against randomly colored dot backgrounds. Background dots were generated by random RGB value selection (0–255) per pixel. Squares displayed pure RGB colors: (1,0,0), (0,1,0), or (0,0,1). Starting from the image center, squares moved with constant velocity in randomly selected directions, reversing direction upon reaching image boundaries. Videos were generated using custom Python scripts.

### Prediction Images

For testing, we prepared two Benham’s top variants: black arcs on a white background and white arcs on a black background (https://doi.org/10.6084/m9.figshare.30215359). Stimulus images were generated using custom Python scripts. Six test conditions were evaluated: stationary, clockwise rotation, and counterclockwise rotation for both variants. Trained networks generated predictions for the 20th frame after processing 19 consecutive input frames.

### Color Analysis

Colors in PredNet-predicted Benham’s top lines were quantified through the following procedure:

1. Line coordinate identification: For black arc stimuli, coordinates were extracted where all RGB values fell below (123, 123, 123); for white arc stimuli, coordinates were extracted where all RGB values exceeded (132, 132, 132). Extracted coordinates were classified into inner and outer circle regions.
2. Color quantification: RGB values were extracted from PredNet output images at the identified line coordinates. Mean values were calculated separately for inner and outer regions, then converted from RGB to HSV format for analysis using a Python library.

## Acknowledgements

The authors would like to thanks Drs. Hidehiko Komatsu and Han Ke for their thoughtful discussion, and also Ms. Tomomi Suginaga for her assistance with the research.

## Author contributions statement

The authors confirm contribution to the paper as follows: study conception and design: Kyohei Ueda, Lana Sinapayen and Eiji Watanabe; data collection and analysis: Kyohei Ueda and Eiji Watanabe; data interpretation of results: Kyohei Ueda and Eiji Watanabe; draft manuscript preparation: Kyohei Ueda and Eiji Watanabe. All authors reviewed the results and approved the final version of the manuscript.

## Statements and Declarations

The authors declared no potential conflicts of interest with respect to the research, authorship, and/or publication of this article.

## Funding statement

This research was conducted with the support of Grants-in-Aid for Scientific Research (A, B, and C) from the Japan Society for the Promotion of Science (to EW), and JST SPRING grant JPMJSP2104 (to KU). Additionally, this research was conducted with the support of Panasonic Automotive Systems Co., Ltd (to EW).

## Ethical consideration

The experimental procedure received approval from the Life Science Research Ethics Committee of the National Institutes of Natural Sciences (Project ID: EC01-078), and informed consent was obtained from all participants.

## AI tools

Large Language Models: The authors received assistance from Claude Sonnet 4 (Anthropic) for language editing and refinement during the revision of the manuscript. All scientific content and conclusions remain the authors’ own work.

## Data availability

The analysis script and data were reposited at Figshare (https://doi.org/10.6084/m9.figshare.30215359).

## Supplemental Tables and Figures

**Table S1.** Human-reported Fechner colors elicited by Benham’s top stimuli in online experiments (n = 98). Participants reported the perceived colors induced by rotating Benham’s tops, and responses were quantified in HSV color space. For each condition, the mean and standard deviation (SD) of Hue, Saturation, and Value are shown separately for the outer and inner arcs. Ba = black-arc variant; Wa = white-arc variant; ccw = counterclockwise rotation; cw = clockwise rotation. Reported hues differ systematically across outer and inner arcs, and their distributions depend on both arc color (black vs. white) and rotation direction.

**Table S2.** Artificial Fechner colors predicted by neural networks trained on natural scene videos (n = 20). PredNet models trained on natural scene dynamics were tested with Benham’s tops, and the predicted colors were quantified in HSV space. For each condition, the mean and standard deviation (SD) of Hue, Saturation, and Value are reported separately for the outer and inner arcs. Ba = black-arc variant; Wa = white-arc variant; ccw = counterclockwise rotation; cw = clockwise rotation; st = static. Predicted hues and saturations emerge predominantly under rotating conditions, whereas static tops produce weak or no color signals.

**Table S3.** Artificial Fechner colors in images predicted by neural networks trained on 3D computer-generated videos (n = 20). PredNet models were trained using computer-generated 3D videos (a walking character in a virtual town) and then tested with Benham’s tops. Predicted colors are summarized in HSV space, with mean and standard deviation (SD) values given separately for outer and inner arcs. Ba = black-arc variant; Wa = white-arc variant; ccw = counterclockwise rotation; cw = clockwise rotation; st = static. Compared with models trained on natural scenes (Table S2), networks trained on structured 3D motion stimuli produced more saturated and consistent color predictions, reflecting the stronger motion–color contingencies encoded during training.

**Table S4.** Artificial Fechner colors in images predicted by neural networks trained on 2D computer-generated videos (n = 20). PredNet models were trained with simplified 2D computer graphics videos consisting of a colored square (red, green, or blue) moving against a dynamic random-dot background. Predicted colors elicited by Benham’s tops are summarized in HSV space, with mean and standard deviation (SD) values shown separately for outer and inner arcs. Ba = black-arc variant; Wa = white-arc variant; ccw = counterclockwise rotation; cw = clockwise rotation; st = static. Depending on the training color, the arcs in the predictions tended to adopt hues similar to the moving square: red-trained models correspond to Figure 6b, green-trained models to Figure 6c, and blue-trained models to Figure 6d. This demonstrates that the learned motion–color contingencies directly shaped the emergence of illusory colors in the model’s outputs.

**Figure S1.** PredNet predictions for Benham’s top variants after training on first-person natural scenes (Ego4D data set). Rose plots depicting the distribution of predicted hues from the neural network model (n = 20 trials). Results are shown for the black-arc variant (top row) and white-arc variant (bottom row) under counterclockwise (ccw, left) and clockwise (cw, right) rotation conditions. Color coding follows the convention established in Figure 2: red histograms represent hues predicted at the outer arc position, and blue histograms represent hues at the inner arc position. Solid lines indicate the mean direction, and dotted lines indicate ±1 standard deviation. The Ego4D numbers of training videos used are as follows: Figure S1a, 3df50de7-7ef7-4938-9cdb-fb2d07751084; S1b, 5c9b85ea-24a9-4bfd-8ac2-4ca6f9090231; S1c, 05df9f6f-90b8-4ed0-81c7-c0e98a730f6f; S1d, 5e11cf07-d5f9-47c0-9e73-2b58028e9a8d; S1e, 22f68c6a-b8dd-44e6-b1fe-d4e619e72f59; S1f, 0963bd23-604b-4e7b-8ae0-19471c4b72ee; S1g, 26192bcd-7240-4d7b-8306-fa4eed0264f2; S1h, 948317e1-7426-4bfa-b25a-a476ee69797f; S1i b6bee905-eddf-4b41-bfe6-d57c9059fe61;, S1j, b5410470-6cb6-43ea-8233-4824ba6a27b0; S1k, c628c1d2-4ce3-4f28-93a4-cc9d3dd9de11; S1l, c25111f0-0c78-4bf8-a16d-57a6c25c2169; S1m, cdfd99eb-88c6-4bc7-8f66-e0318216feab; 1n, d41f6dac-369a-4fed-abed-2423b28bb8f8; S1o, d437be8d-6f9c-4ce8-b863-a7f21c39d1df; S1p, e9725499-415a-490c-a1c7-6089030c958a. The first 40,000 frames of each video were used for training.

**Figure S2.** Visualization of node-block activations in Layer 3 Convolution LSTM during Benham’s top stimulation. In this experiment, 40,000 frames were randomly extracted from the Moments in Time dataset (http://moments.csail.mit.edu/, a massive dataset of collected 3-second videos) and used as training videos. ASC robustly emerged even when using completely different types of videos. At the time step after inputting 20 image sequences of the Benham’s top test stimuli, the activation patterns of the hidden units of PredNet layers were visualized. Visualization was performed using a self-made converter program using TensorBoard in TensorFlow (https://doi.org/10.6084/m9.figshare.30215359). As a reference for the discussion, we have provided the activation patterns of one particular unit out of 181 hidden units. (a) A representative predicted images obtained during the inference phase when Benham’s tops (ccw) were presented as novel inputs. (b) Activation patterns of node blocks in Layer 3 of the Conv LSTM module when the black-arc variant of Benham’s top was rotated counterclockwise. Each small square represents a block composed of multiple nodes, revealing clusters of units with related spatial selectivity. Blue boxes highlight blocks that responded strongly to arc-related features. (c) Enlarged views of the arc-related node blocks from panel b (orange boxes, outer arc-related; blue boxes, inner arc related). These activations consistently exhibit a composite structure in which the arc is accompanied by an attached quarter-sector (¼ fan-shaped segment). This suggests that the internal representation encodes not only the arc itself but also adjacent contextual features, reflecting how the model decomposes dynamic stimuli into structured components.

## Notes

https://doi.org/10.6084/m9.figshare.30215359

### Competing Interest Statement

The authors have declared no competing interest.

