## Supplemental Tables and Figures for "Predictive Networks Generate Motion-Induced Color Illusions"

#### Table S1

| Table S1 | | outer arc hue mean | outer arc hue SD | inner arc hue mean | inner arc hue SD | | outer arc saturation mean | outer arc saturation SD | inner arc saturation mean | inner arc saturation SD | | outer arc value mean | outer arc value SD | inner arc value mean | inner arc value SD |
|---|---|---|---|---|---|---|---|---|---|---|---|---|---|---|---|
| | Ba ccw | 260.3 | 63.0 | 53.3 | 73.4 | | 47.4 | 29.7 | 40.5 | 34.3 | | 88.0 | 19.1 | 96.1 | 9.9 |
| | Ba cw | 63.2 | 68.7 | 268.5 | 57.7 | | 37.4 | 33.2 | 44.3 | 28.1 | | 94.4 | 14.0 | 90.5 | 17.8 |
| | Wa ccw | 72.1 | 65.7 | 272.8 | 70.9 | | 69.0 | 28.8 | 65.4 | 35.3 | | 80.2 | 26.4 | 67.0 | 37.9 |
| | Wa cw | 282.6 | 93.9 | 59.2 | 75.9 | | 74.2 | 32.4 | 67.3 | 26.6 | | 68.2 | 35.2 | 78.7 | 27.5 |

#### Table S2

| Table S2 | | outer arc hue mean | outer arc hue SD | inner arc hue mean | inner arc hue SD | | outer arc saturation mean | outer arc saturation SD | inner arc saturation mean | inner arc saturation SD | | outer arc value mean | outer arc value SD | inner arc value mean | inner arc value SD |
|---|---|---|---|---|---|---|---|---|---|---|---|---|---|---|---|
| | Ba ccw | 201.3 | 75.6 | 27.3 | 60.3 | | 12.9 | 10.2 | 12.0 | 6.4 | | 70.4 | 4.7 | 63.5 | 3.9 |
| | Ba cw | 29.8 | 60.2 | 193.5 | 78.5 | | 11.8 | 6.0 | 13.4 | 10.8 | | 62.3 | 3.7 | 70.6 | 4.6 |
| | Ba st | 53.8 | 60.6 | 59.2 | 53.3 | | 0.7 | 0.6 | 0.6 | 0.6 | | 58.2 | 1.2 | 59.1 | 1.1 |
| | Wa ccw | 19.5 | 46.6 | 261.7 | 74.8 | | 25.4 | 10.0 | 7.5 | 4.3 | | 48.8 | 6.8 | 46.8 | 3.3 |
| | Wa cw | 255.8 | 64.7 | 18.1 | 44.5 | | 10.4 | 5.6 | 20.9 | 9.5 | | 42.2 | 3.7 | 54.2 | 5.9 |
| | Wa st | 284.4 | 40.0 | 264.3 | 35.7 | | 2.7 | 0.8 | 1.7 | 0.6 | | 40.2 | 1.2 | 45.6 | 1.1 |

#### Table S3

| Table S3 | | outer arc hue mean | outer arc hue SD | inner arc hue mean | inner arc hue SD | | outer arc saturation mean | outer arc saturation SD | inner arc saturation mean | inner arc saturation SD | | outer arc value mean | outer arc value SD | inner arc value mean | inner arc value SD |
|---|---|---|---|---|---|---|---|---|---|---|---|---|---|---|---|
| | Ba ccw | 208.3 | 17.0 | 308.0 | 20.0 | | 34.1 | 11.8 | 27.2 | 10.5 | | 51.8 | 3.8 | 31.3 | 3.7 |
| | Ba cw | 301.1 | 19.5 | 209.6 | 18.2 | | 30.2 | 12.7 | 30.0 | 11.7 | | 27.2 | 3.4 | 53.4 | 4.1 |
| | Ba st | 94.4 | 42.0 | 71.5 | 43.1 | | 1.3 | 0.7 | 1.9 | 1.0 | | 36.3 | 0.3 | 37.3 | 0.3 |
| | Wa ccw | 335.8 | 12.7 | 225.4 | 26.8 | | 31.0 | 7.6 | 11.3 | 7.4 | | 60.0 | 6.8 | 63.5 | 2.9 |
| | Wa cw | 223.2 | 25.0 | 333.5 | 13.1 | | 14.8 | 9.2 | 26.9 | 6.7 | | 64.8 | 3.7 | 63.2 | 6.1 |
| | Wa st | 242.1 | 82.0 | 222.3 | 48.1 | | 0.7 | 0.2 | 0.5 | 0.2 | | 63.0 | 0.4 | 64.7 | 0.2 |

#### Table S4

| Table S4a (red) | | outer arc hue mean | outer arc hue SD | inner arc hue mean | inner arc hue SD | | outer arc saturation mean | outer arc saturation SD | inner arc saturation mean | inner arc saturation SD | | outer arc value mean | outer arc value SD | inner arc value mean | inner arc value SD |
|---|---|---|---|---|---|---|---|---|---|---|---|---|---|---|---|
| | Ba ccw | 202.6 | 51.4 | 327.7 | 57.8 | | 53.3 | 18.4 | 43.1 | 18.8 | | 58.6 | 11.0 | 58.8 | 11.8 |
| | Ba cw | 330.9 | 53.4 | 203.1 | 52.1 | | 44.8 | 18.7 | 52.8 | 18.7 | | 58.8 | 11.8 | 60.0 | 11.2 |
| | Ba st | 69.6 | 102.4 | 60.9 | 94.5 | | 10.0 | 7.5 | 9.7 | 8.0 | | 31.6 | 1.8 | 32.4 | 1.8 |
| | Wa ccw | 346.4 | 20.2 | 280.1 | 71.9 | | 68.4 | 13.2 | 26.2 | 14.1 | | 78.8 | 13.5 | 75.6 | 7.0 |
| | Wa cw | 265.4 | 80.3 | 345.6 | 20.5 | | 31.0 | 15.7 | 57.8 | 11.5 | | 73.6 | 7.3 | 80.2 | 12.2 |
| | Wa st | 352.6 | 78.1 | 321.9 | 70.0 | | 2.0 | 1.0 | 2.2 | 1.3 | | 70.9 | 0.7 | 72.2 | 0.6 |

| Table S4b (green) | | outer arc hue mean | outer arc hue SD | inner arc hue mean | inner arc hue SD | | outer arc saturation mean | outer arc saturation SD | inner arc saturation mean | inner arc saturation SD | | outer arc value mean | outer arc value SD | inner arc value mean | inner arc value SD |
|---|---|---|---|---|---|---|---|---|---|---|---|---|---|---|---|
| | Ba ccw | 303.9 | 48.9 | 96.5 | 59.8 | | 45.7 | 19.1 | 43.2 | 21.7 | | 46.5 | 14.7 | 49.7 | 10.1 |
| | Ba cw | 98.6 | 54.7 | 305.0 | 49.6 | | 43.4 | 21.5 | 48.0 | 19.3 | | 50.7 | 10.8 | 46.4 | 14.3 |
| | Ba st | 188.1 | 44.2 | 182.1 | 59.0 | | 9.5 | 6.8 | 9.3 | 6.3 | | 33.3 | 2.8 | 33.1 | 2.2 |
| | Wa ccw | 118.7 | 4.6 | 118.9 | 82.9 | | 75.3 | 9.6 | 17.3 | 9.3 | | 81.7 | 11.7 | 70.0 | 9.9 |
| | Wa cw | 109.5 | 79.0 | 119.9 | 6.1 | | 21.9 | 10.7 | 65.1 | 9.0 | | 68.6 | 12.0 | 81.1 | 9.8 |
| | Wa st | 13.4 | 94.2 | 126.0 | 71.0 | | 1.1 | 0.6 | 1.8 | 1.2 | | 69.9 | 0.8 | 71.4 | 0.5 |

| Table S4c (blue) | | outer arc hue mean | outer arc hue SD | inner arc hue mean | inner arc hue SD | | outer arc saturation mean | outer arc saturation SD | inner arc saturation mean | inner arc saturation SD | | outer arc value mean | outer arc value SD | inner arc value mean | inner arc value SD |
|---|---|---|---|---|---|---|---|---|---|---|---|---|---|---|---|
| | Ba ccw | 189.6 | 86.9 | 254.3 | 104.3 | | 30.2 | 14.8 | 21.7 | 11.7 | | 53.5 | 9.7 | 50.2 | 7.4 |
| | Ba cw | 222.7 | 100.7 | 205.8 | 86.3 | | 23.0 | 11.9 | 31.1 | 15.2 | | 51.0 | 7.4 | 52.6 | 10.1 |
| | Ba st | 87.9 | 109.3 | 95.4 | 88.4 | | 12.3 | 13.8 | 17.7 | 23.4 | | 32.3 | 3.3 | 32.1 | 2.2 |
| | Wa ccw | 235.1 | 43.6 | 197.5 | 98.2 | | 54.9 | 20.3 | 19.8 | 10.2 | | 65.8 | 17.2 | 69.9 | 7.9 |
| | Wa cw | 177.1 | 112.4 | 235.9 | 40.6 | | 22.0 | 9.8 | 48.8 | 16.3 | | 70.4 | 9.9 | 69.8 | 16.6 |
| | Wa st | 314.6 | 70.9 | 218.5 | 85.9 | | 2.7 | 1.9 | 4.5 | 5.0 | | 69.8 | 1.0 | 70.4 | 1.3 |

**Figure S1a**

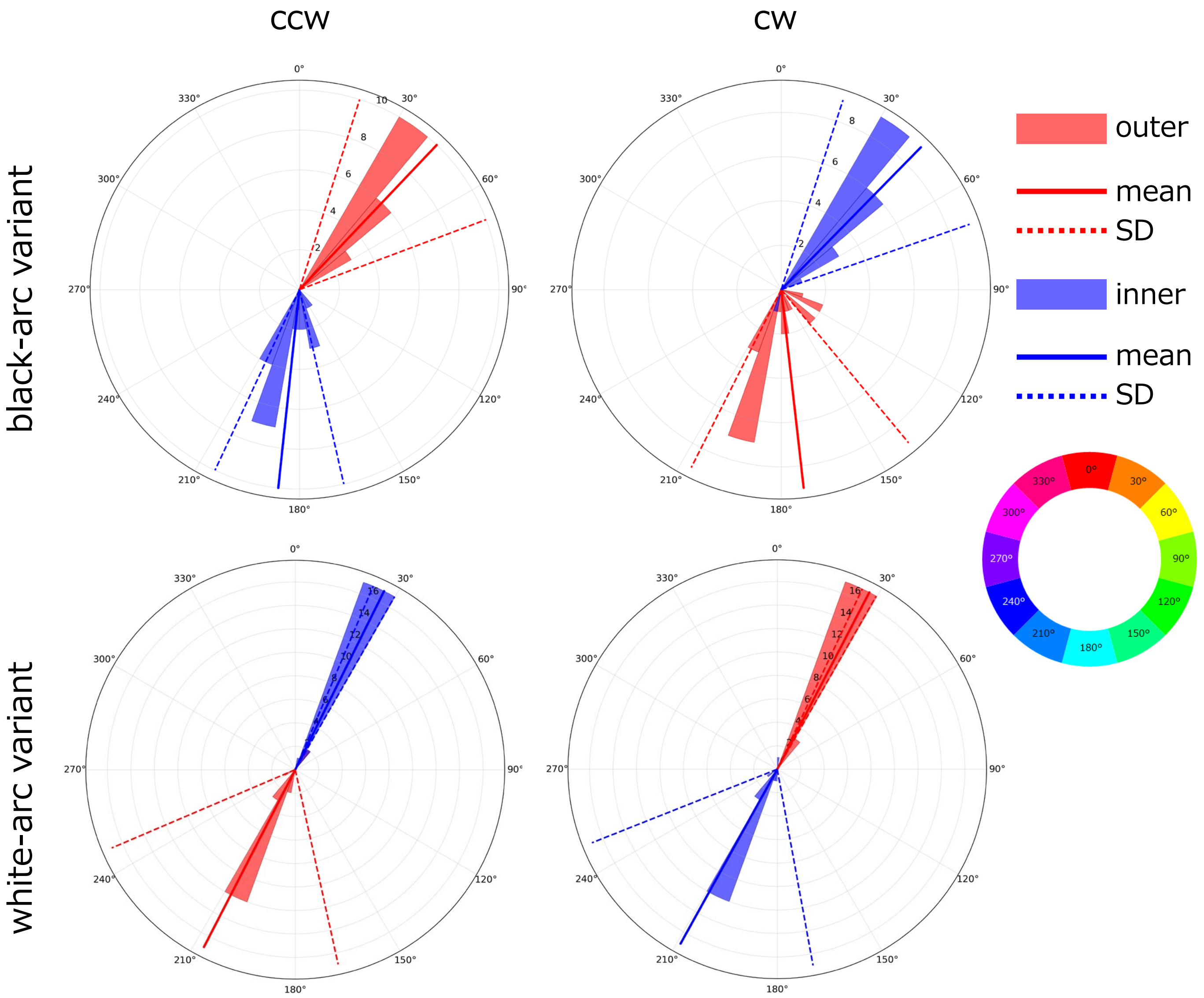


Video No. 3df50de7-7ef7-4938-9cdb-fb2d07751084

**Figure S1b**

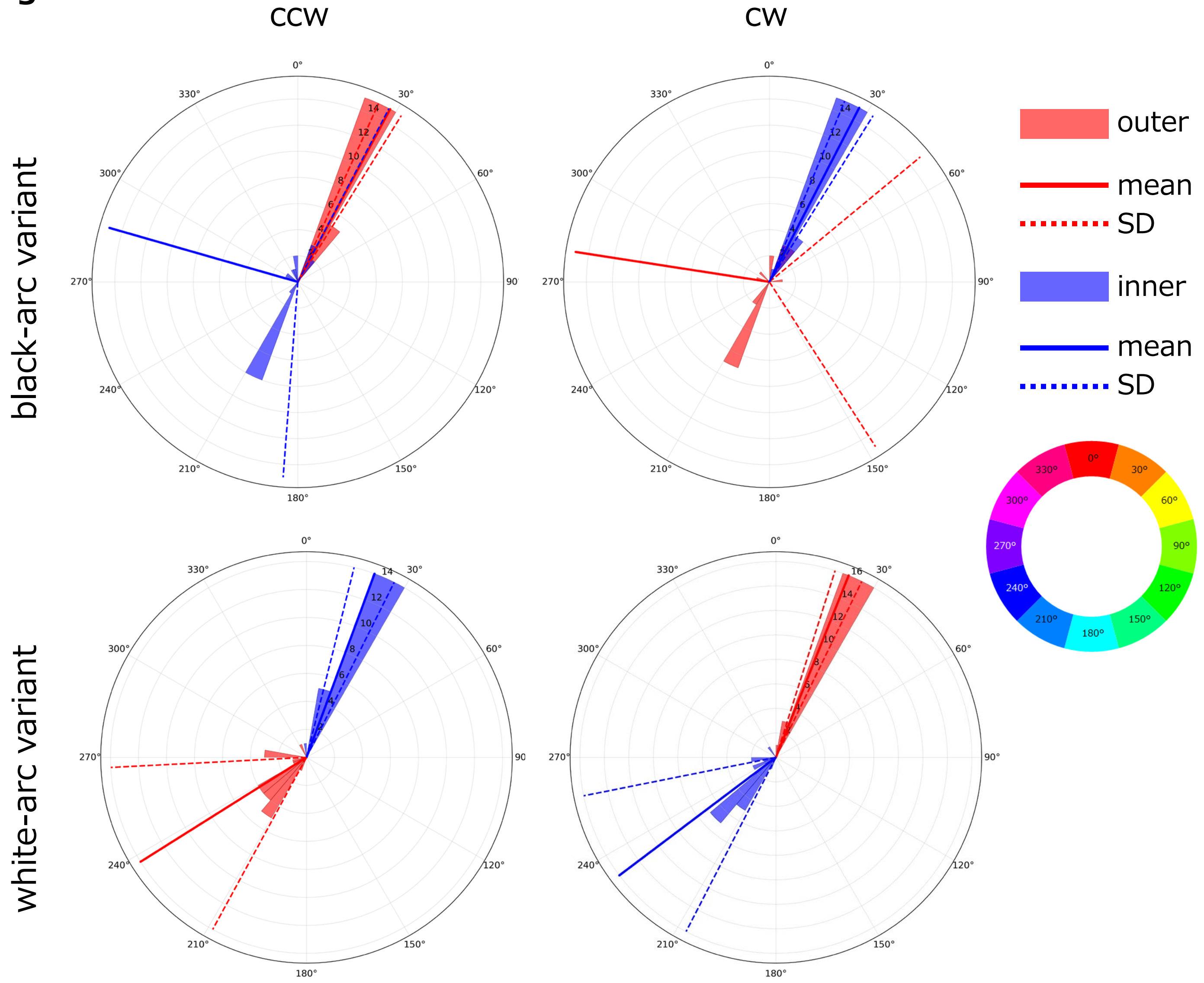


Video No. 5c9b85ea-24a9-4bfd-8ac2-4ca6f9090231

**Figure S1c**

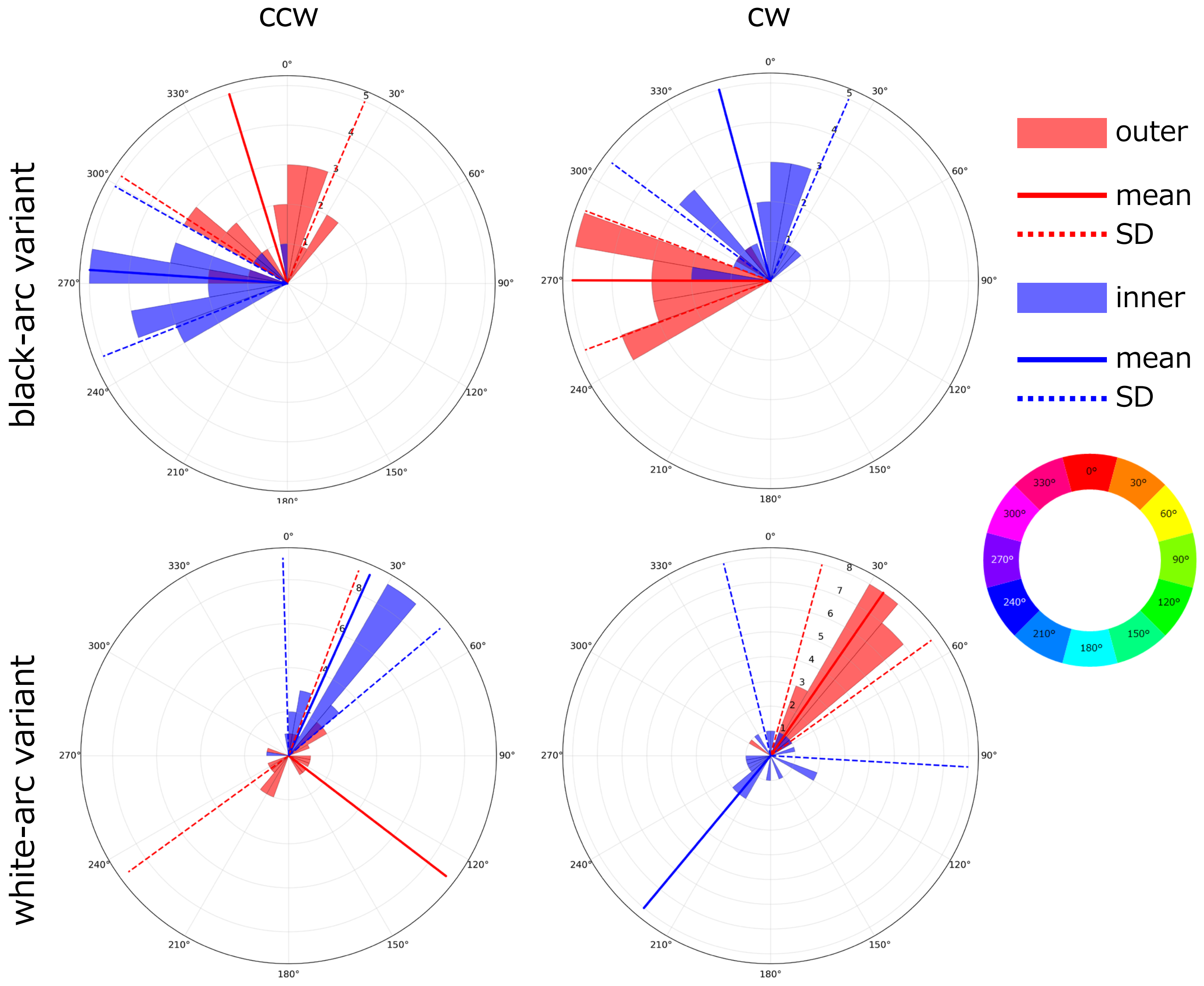


Video No. 05df9f6f-90b8-4ed0-81c7-c0e98a730f6f

#### Figure S1d

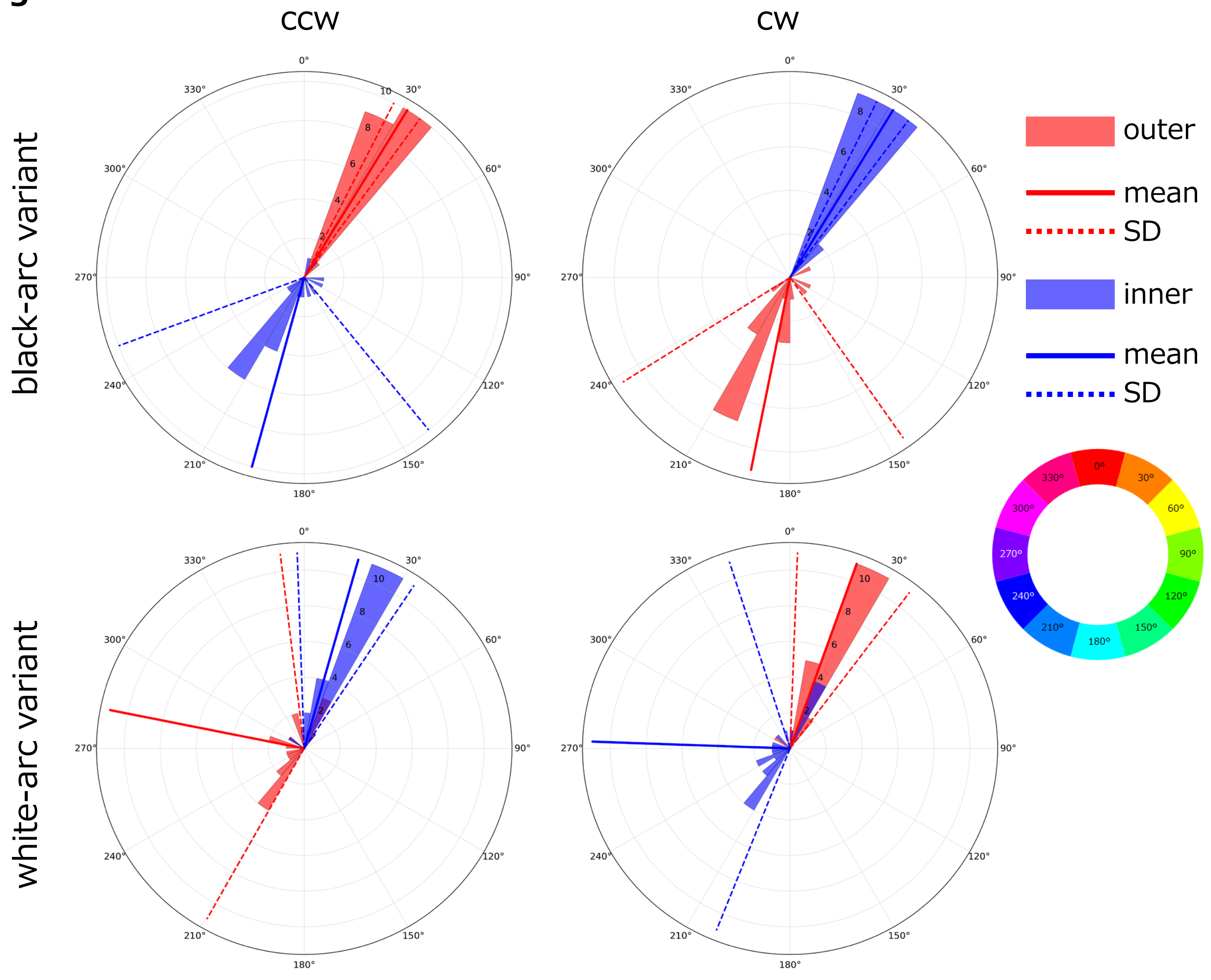


Video No. 5e11cf07-d5f9-47c0-9e73-2b58028e9a8d

#### Figure S1e

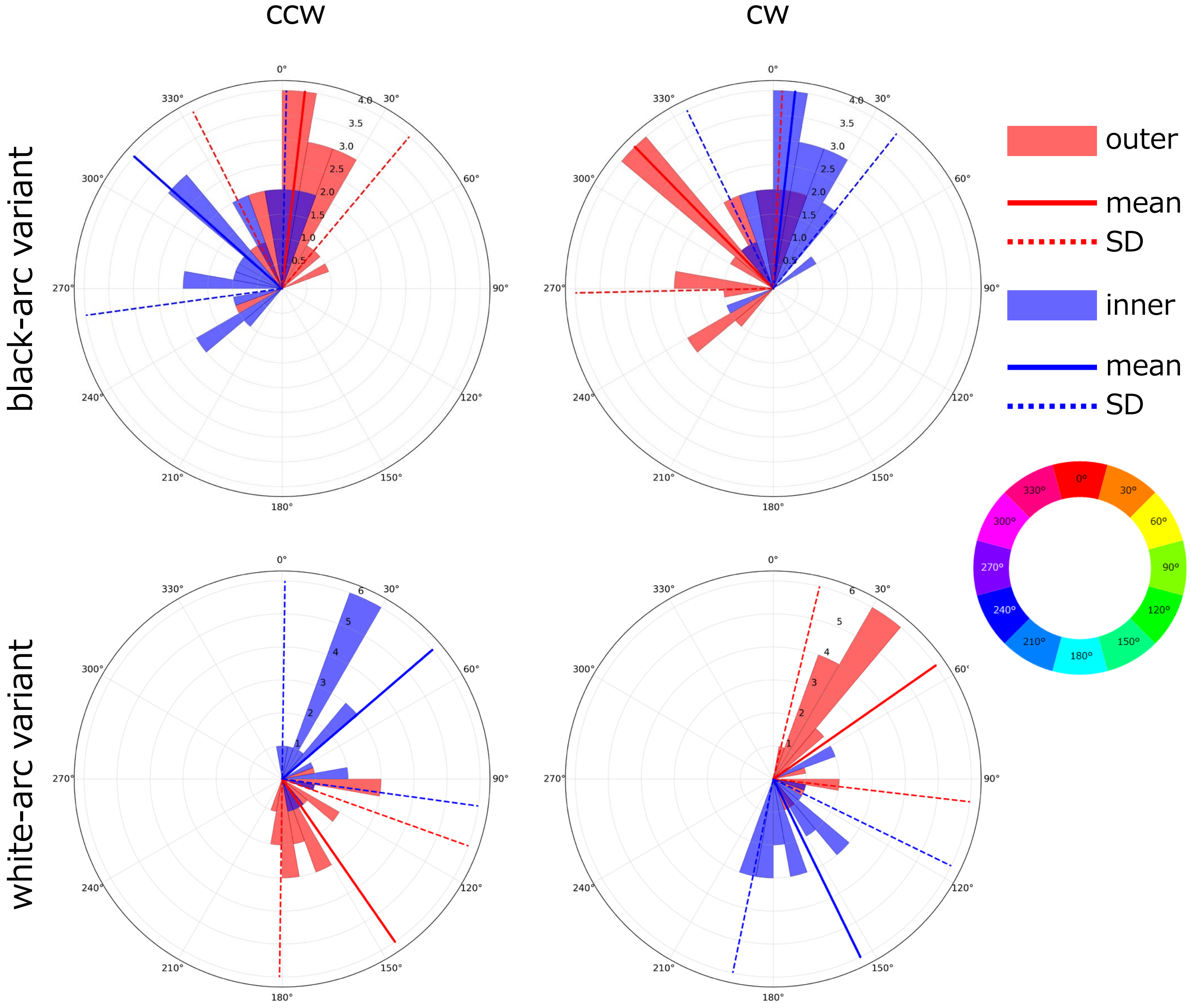


Video No. 22f68c6a-b8dd-44e6-b1fe-d4e619e72f59

**Figure S1f**

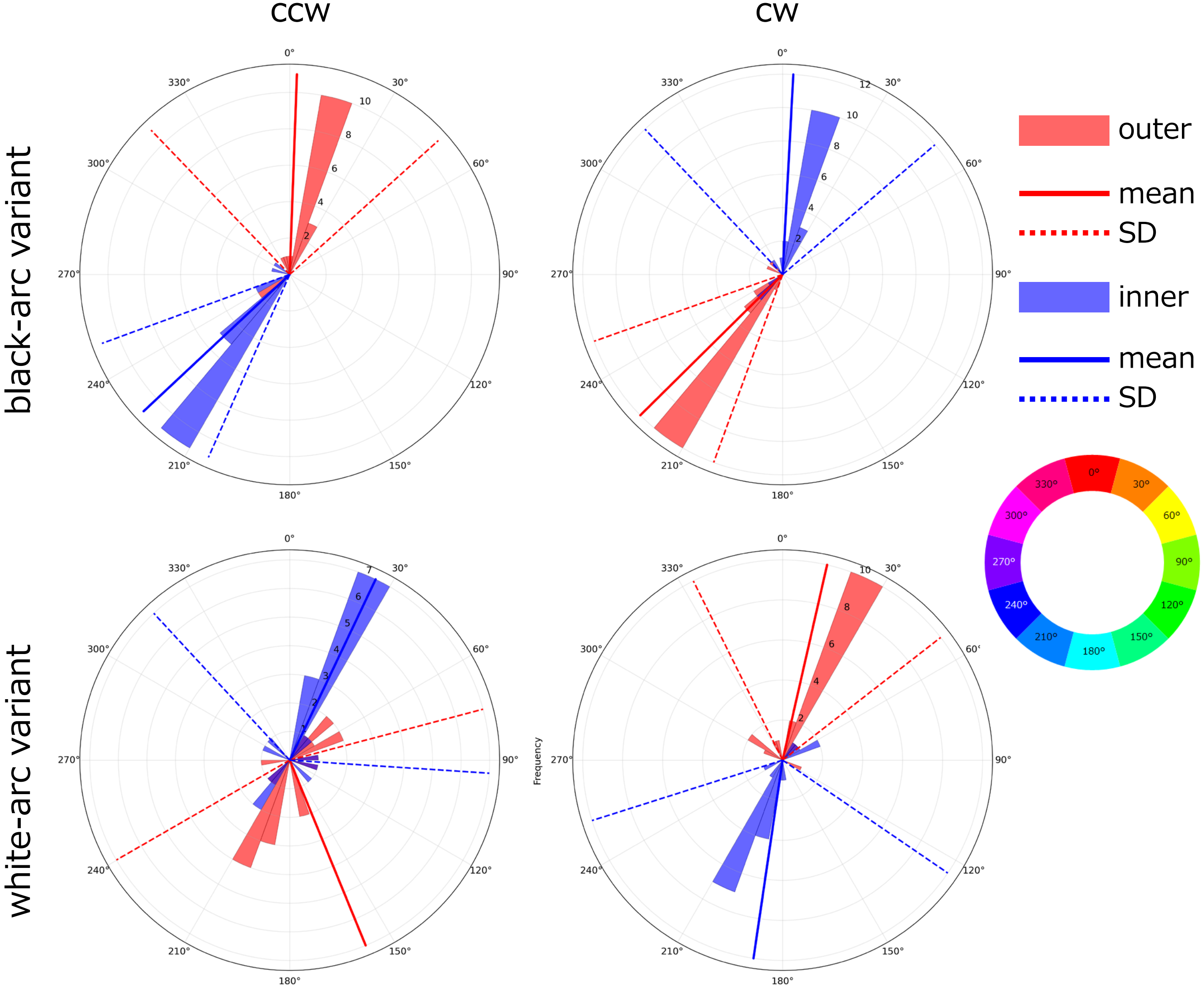


Video No. 0963bd23-604b-4e7b-8ae0-19471c4b72ee

#### Figure S1g

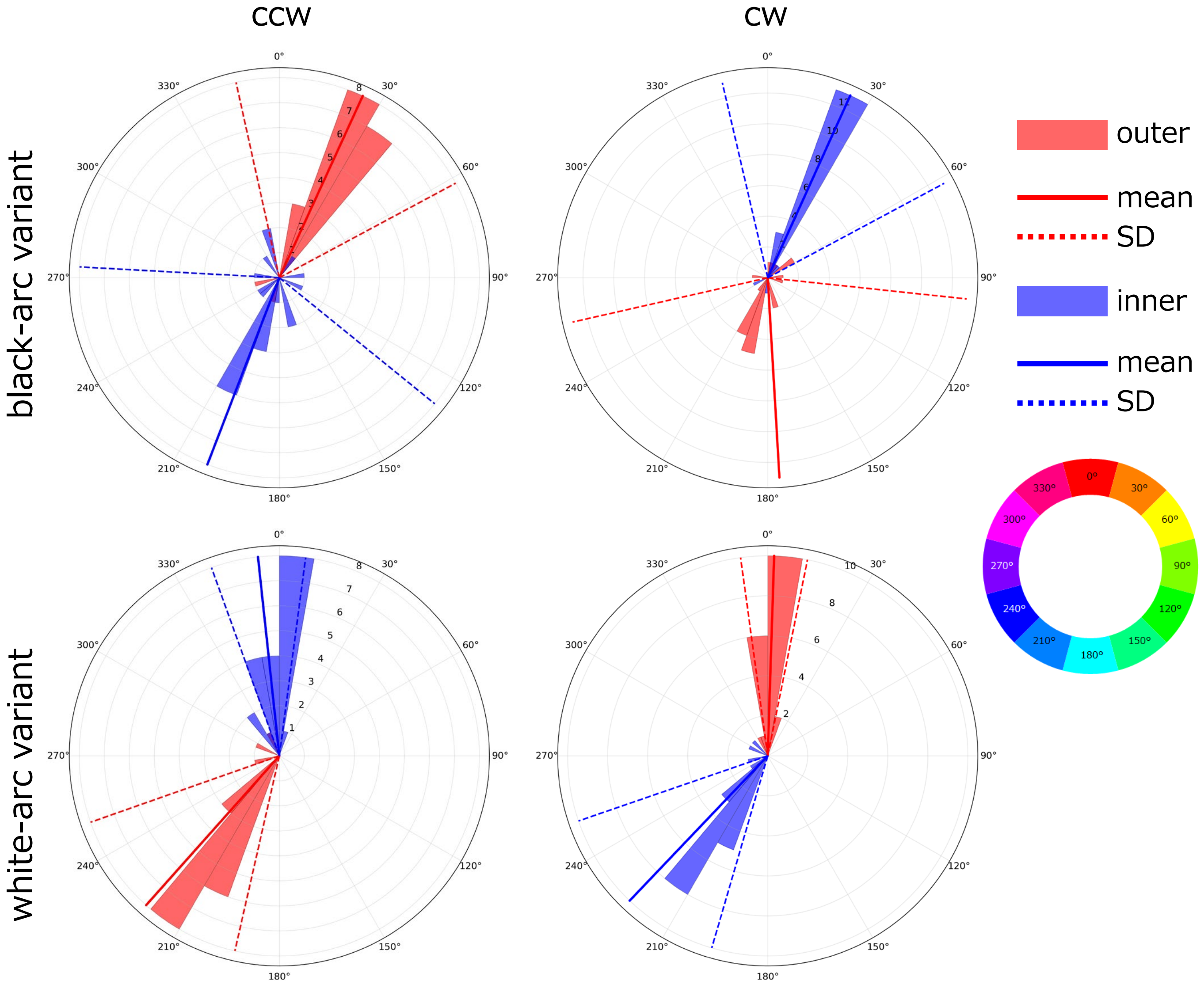


Video No. 26192bcd-7240-4d7b-8306-fa4eed0264f2

**Figure S1h**

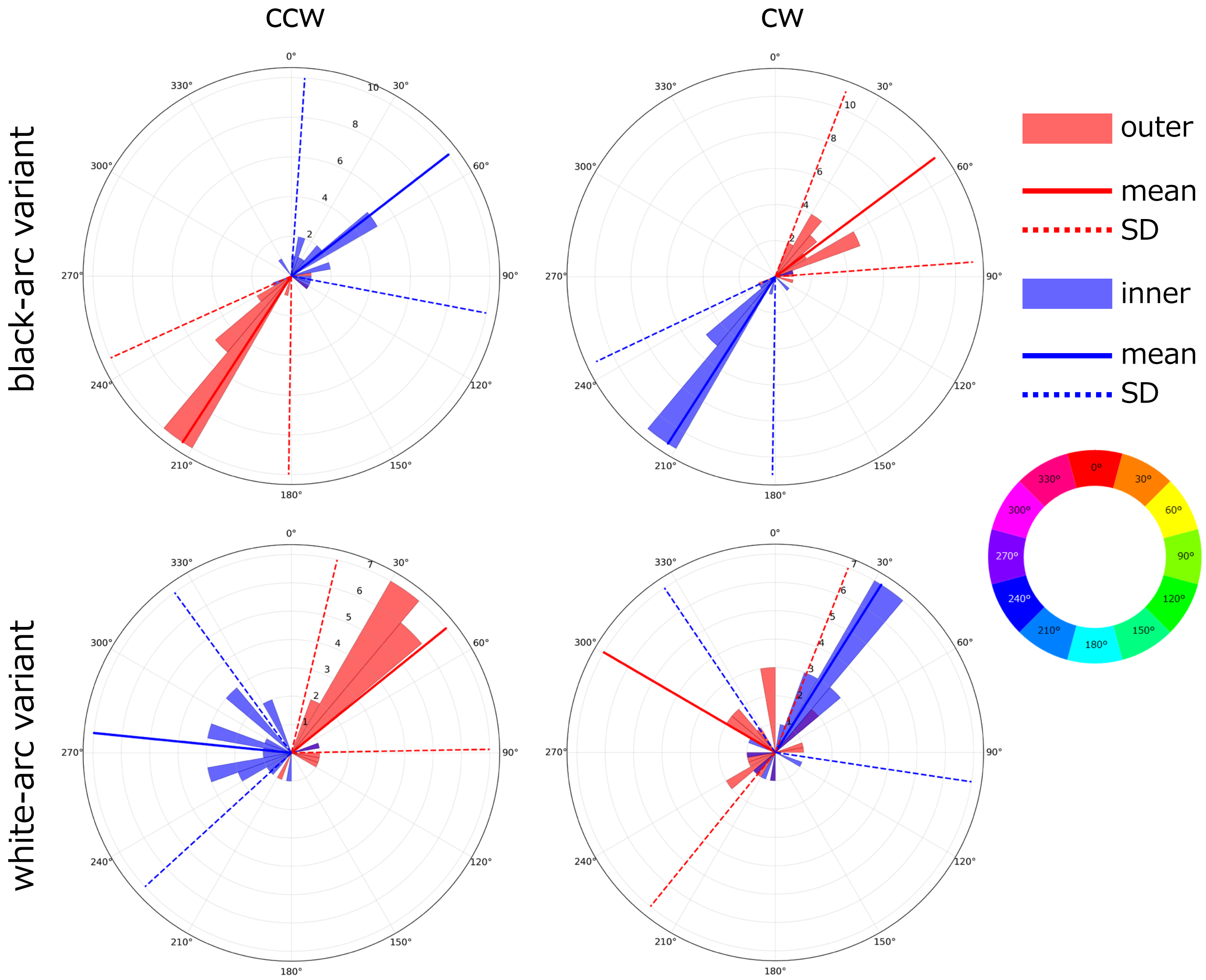


Video No. 948317e1-7426-4bfa-b25a-a476ee69797f

#### Figure S1i

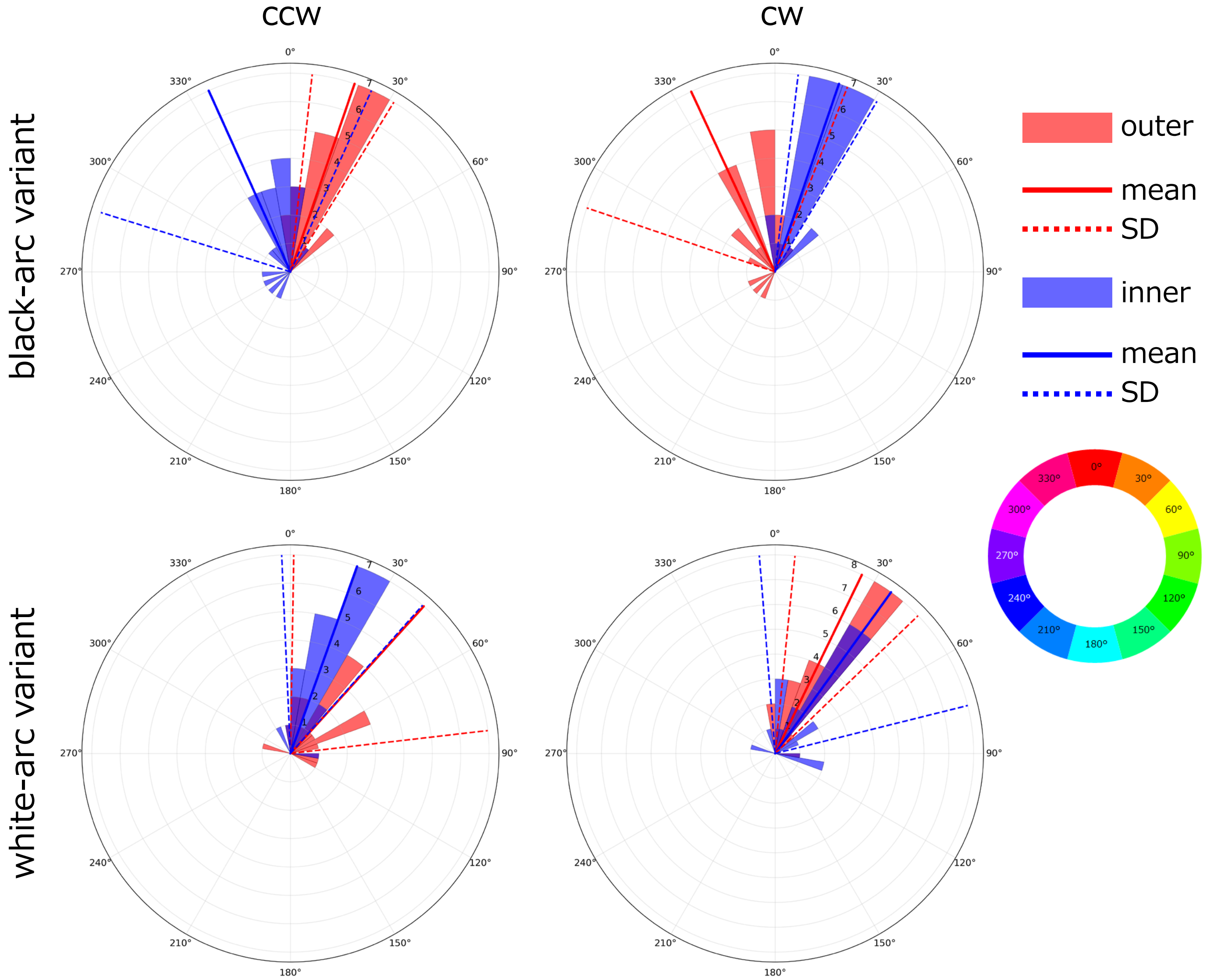


Video No. b6bee905-eddf-4b41-bfe6-d57c9059fe61

**Figure S1j**

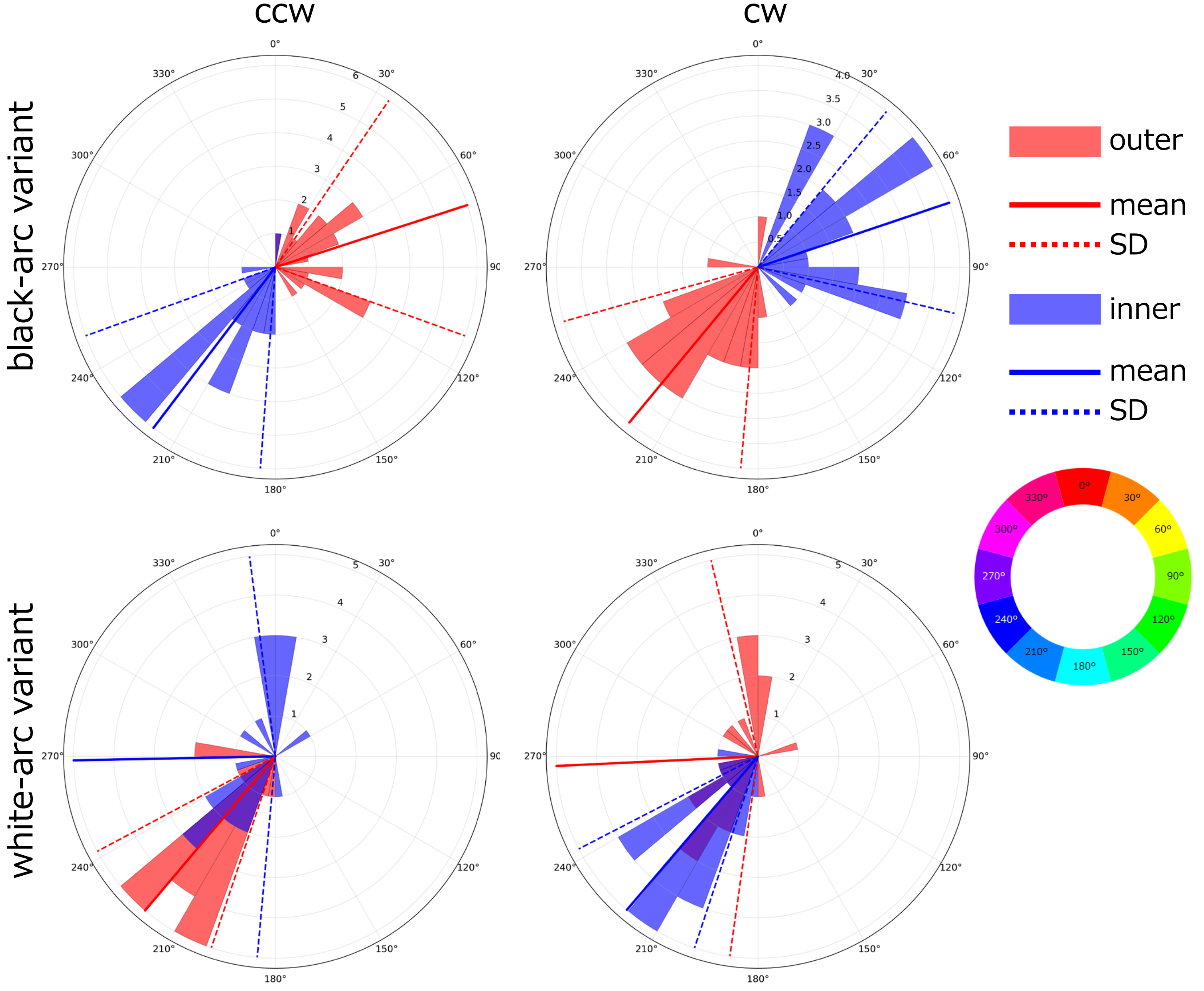


Video No. b5410470-6cb6-43ea-8233-4824ba6a27b0

**Figure S1k**

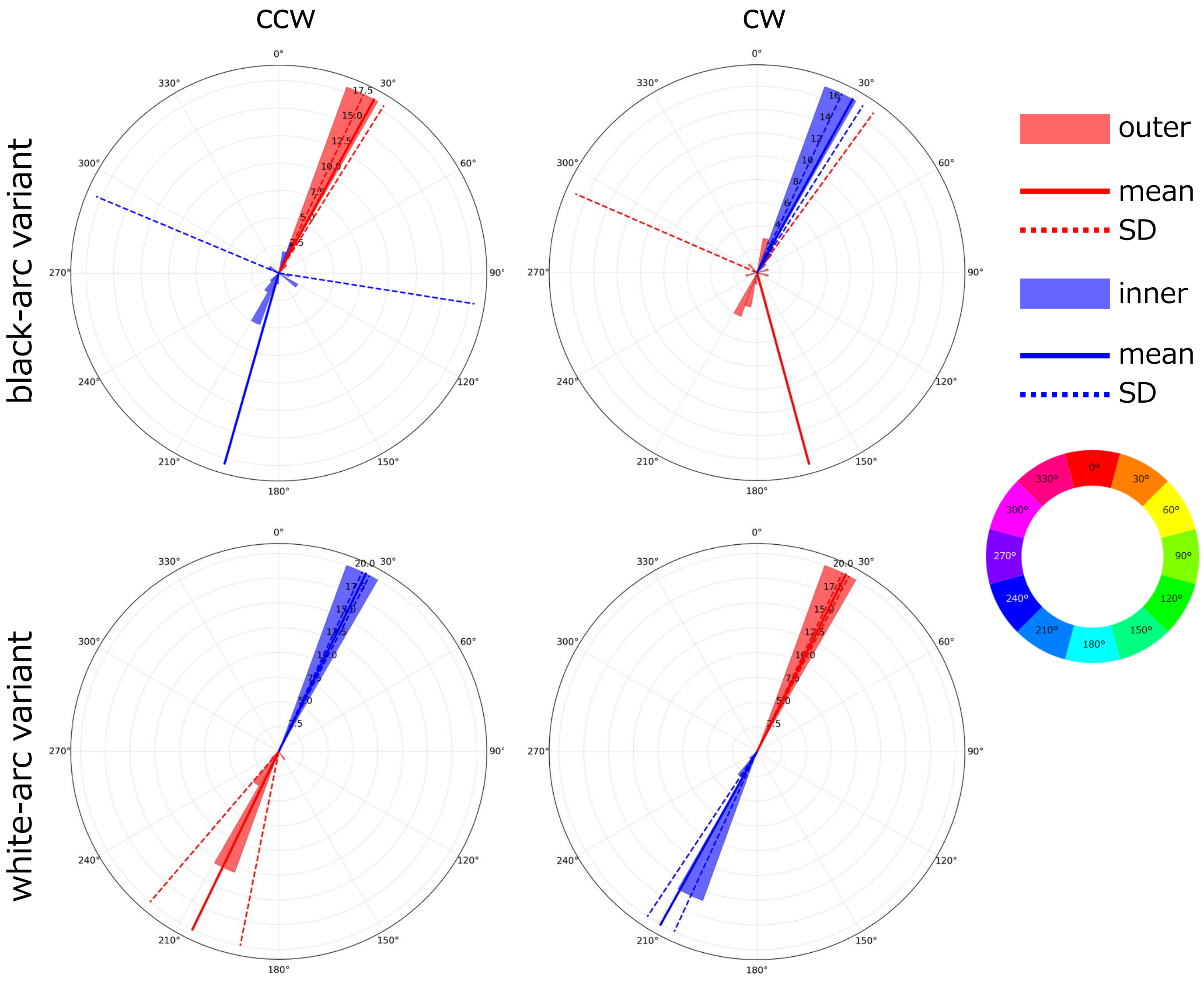


Video No. c628c1d2-4ce3-4f28-93a4-cc9d3dd9de11

**Figure S1l**

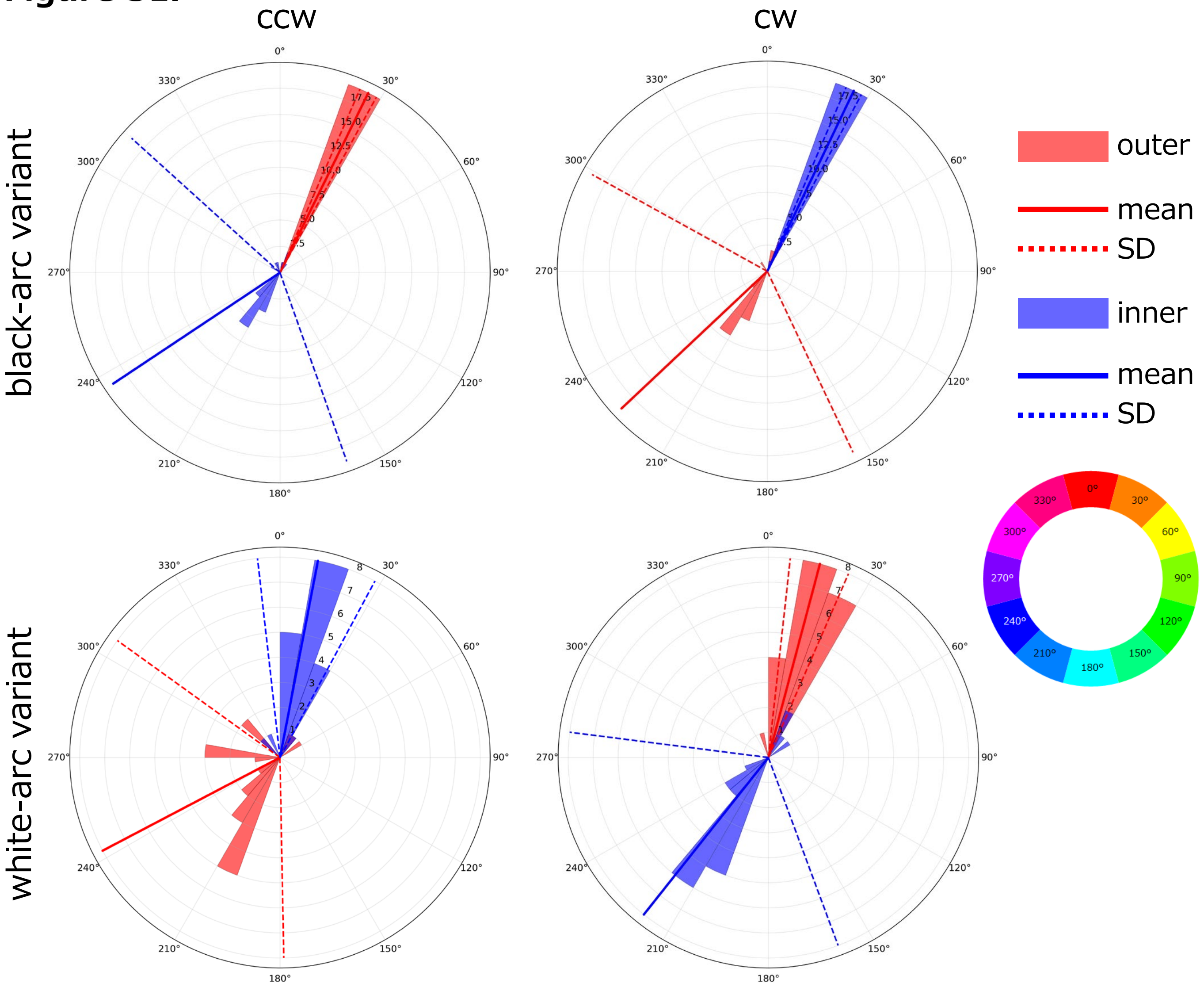


Video No. c25111f0-0c78-4bf8-a16d-57a6c25c2169

#### Figure S1m

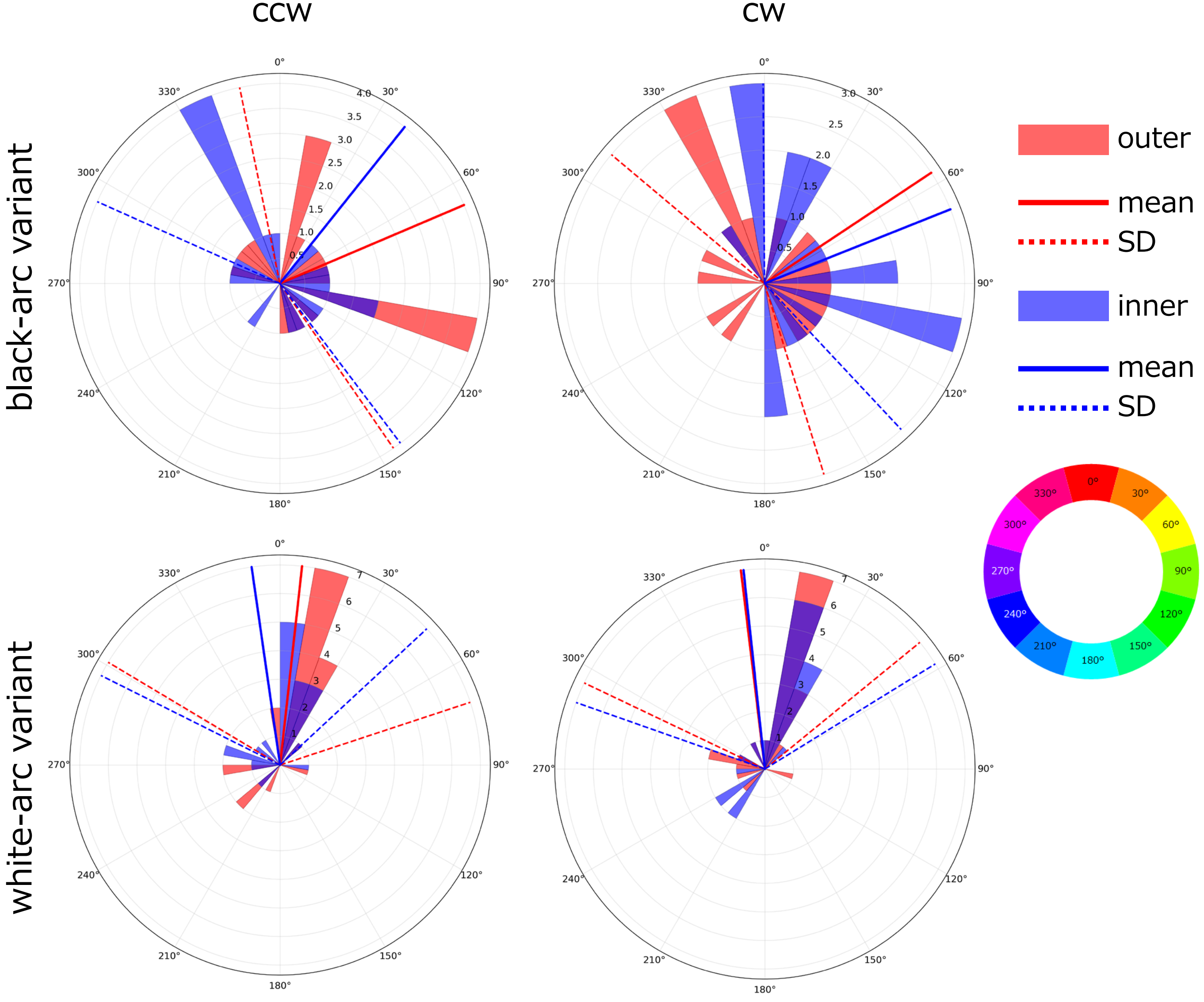


Video No. cdfd99eb-88c6-4bc7-8f66-e0318216feab

#### Figure S1n

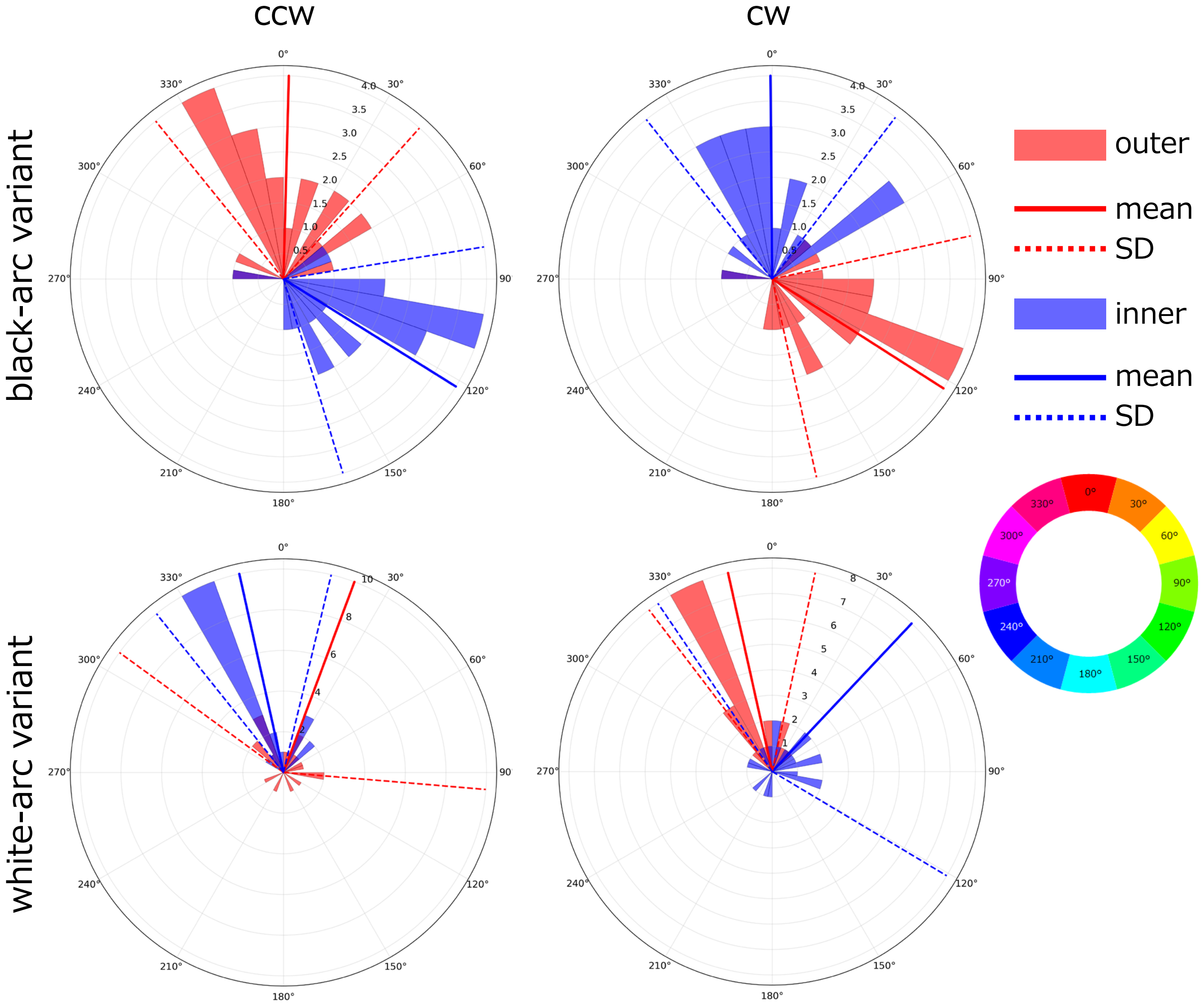


Video No. d41f6dac-369a-4fed-abed-2423b28bb8f8

### Figure S1o

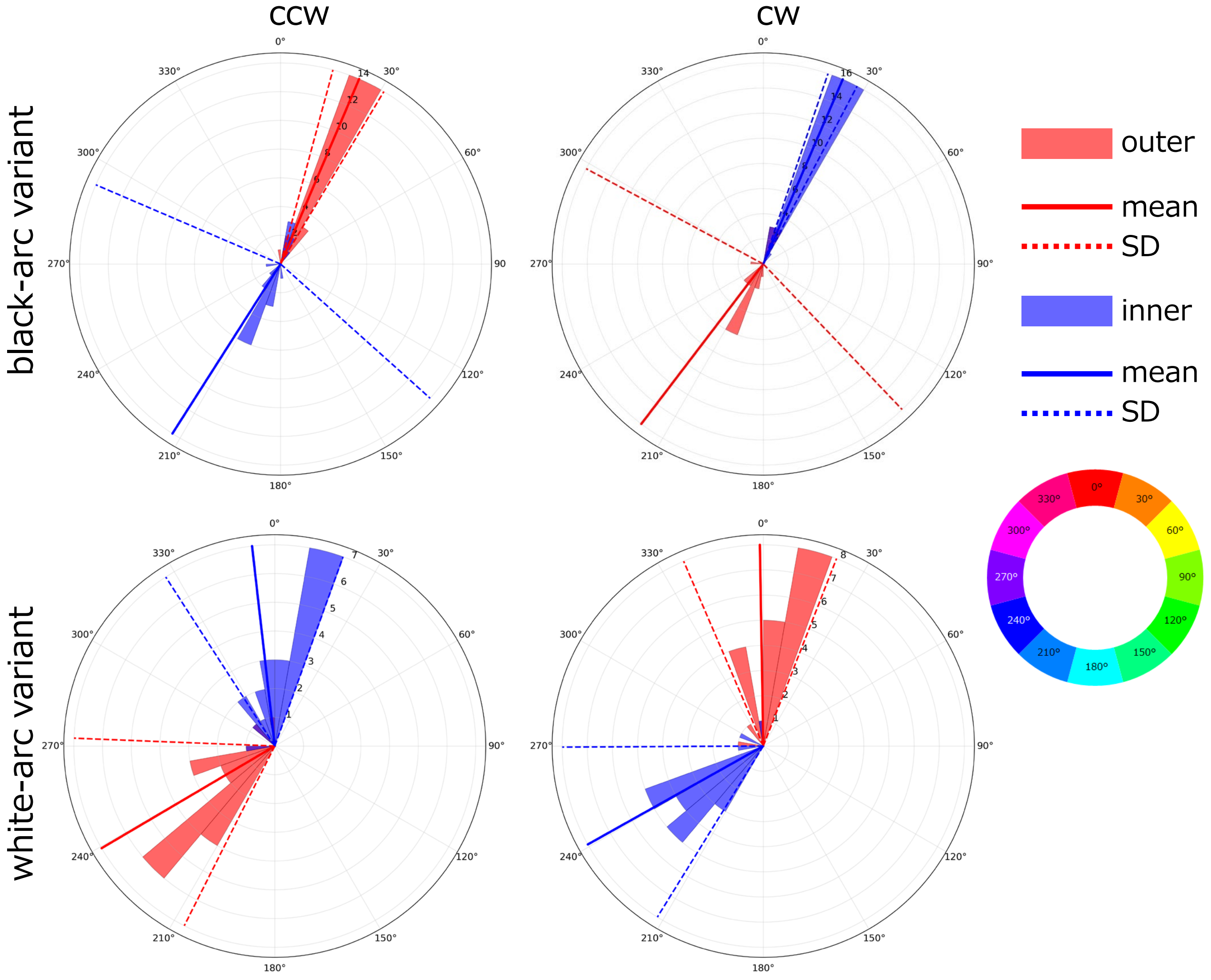


Video No. d437be8d-6f9c-4ce8-b863-a7f21c39d1df

#### Figure S1p

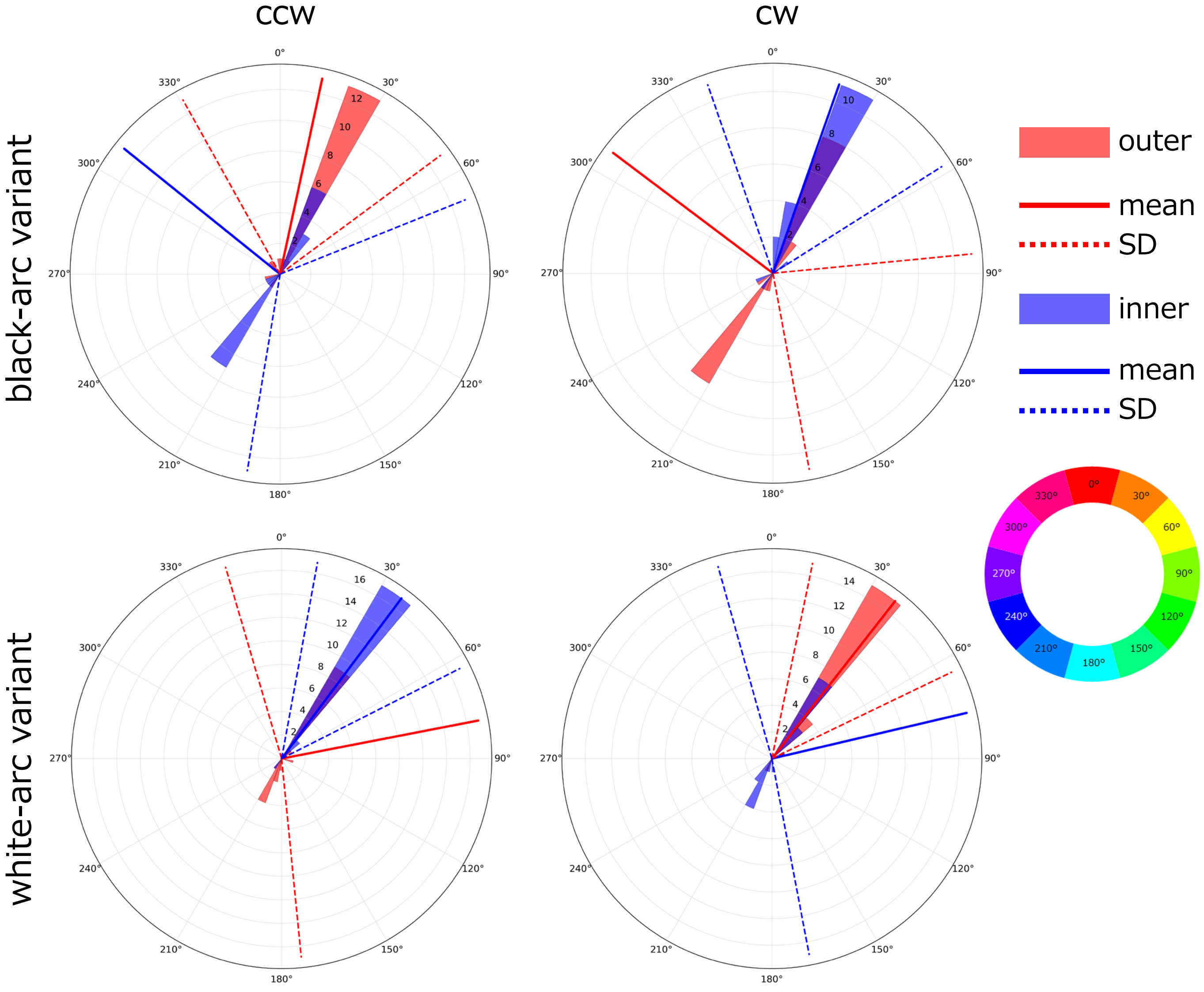


Video No. e9725499-415a-490c-a1c7-6089030c958a

#### Figure S2

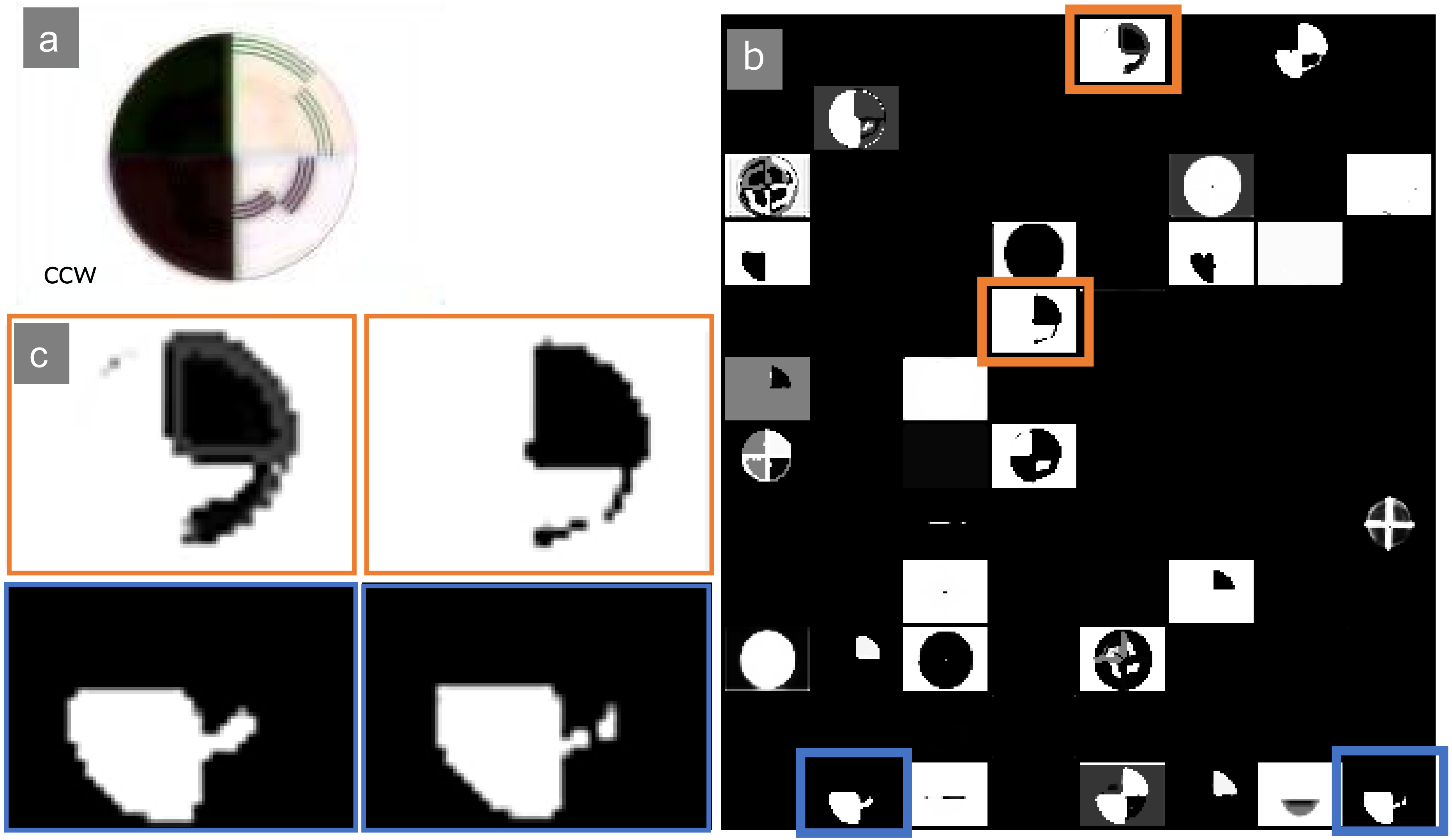
